# ModulOmics: Integrating Multi-Omics Data to Identify Cancer Driver Modules

**DOI:** 10.1101/288399

**Authors:** Dana Silverbush, Simona Cristea, Gali Yanovich, Tamar Geiger, Niko Beerenwinkel, Roded Sharan

## Abstract

The identification of molecular pathways driving cancer progression is a fundamental unsolved problem in tumorigenesis, which can substantially further our understanding of cancer mechanisms and inform the development of targeted therapies. Most current approaches to address this problem use primarily somatic mutations, not fully exploiting additional layers of biological information. Here, we describe ModulOmics, a method to *de novo* identify cancer driver pathways, or modules, by integrating multiple data types (protein-protein interactions, mutual exclusivity of mutations or copy number alterations, transcriptional co-regulation, and RNA co-expression) into a single probabilistic model. To efficiently search the exponential space of candidate modules, ModulOmics employs a two-step optimization procedure that combines integer linear programming with stochastic search. Across several cancer types, ModulOmics identifies highly functionally connected modules enriched with cancer driver genes, outperforming state-of-the-art methods. For breast cancer subtypes, the inferred modules recapitulate known molecular mechanisms and suggest novel subtype-specific functionalities. These findings are supported by an independent patient cohort, as well as independent proteomic and phosphoproteomic datasets.

## Introduction

Rapid advancements in sequencing technologies led to an unprecedented increase in the generation and availability of high-resolution DNA, RNA, and protein cancer data. These large datasets are analyzed with mathematical and computational tools, unveiling mechanistic and predictive insights into cancer progression and treatment [5, 7, 8]. Key to achieving these goals is the identification of molecular alterations that drive tumorigenesis, or drivers, such as single nucleotide variants (SNVs), copy number alterations (CNAs), changes in the transcriptional activity of genes, or changes in protein concentration. Groups of such functionally connected genetic alterations, also termed cancer driver modules or pathways, activate mechanisms that drive tumorigenesis and gradually contribute to triggering the hallmarks of cancer, conferring fitness advantages to the tumors [54, 22]. Driver module elucidation can further our understanding of cancer initiation and progression, as well as inform the development of targeted therapies.

It has been observed that members of cancer pathways often display specific alteration patterns across tumor samples, most notably co-occurrence and mutual exclusivity [12, 11, 30, 2]. Finding groups of mutually exclusive genes is an efficient way to identify cancer modules fulfilling the same biological function, since, once a single member of the group is altered, the tumor gains a significant selective advantage, and the fitness of the tumor is not expected to increase with the alteration of additional group members. However, most existing mutual exclusivity methods rely only on DNA-level data, particularly SNVs, failing to fully exploit the complex interactions involving RNA or protein molecules potentially driving tumorigenesis [14, 19]. To this end, data integration strategies have the potential to unravel previously unknown cancer driver modules.

An additional important data source for identifying interactions among cancer drivers is protein-protein interaction (PPI) networks, cataloged in databases such as HIPPIE [43], STRING [48], or BioGRID [46]. Studies that exploit this data source include HotNet2 [29], which uses PPI networks of genetic alterations to identify significantly altered subnetworks connecting recurrently mutated genes; EnrichNet [20], which identifies functional gene sets based on PPI proximity calculated similarly to HotNet2, MEMo [11], which identifies mutually exclusive gene sets on the basis of PPI-filtered pairwise connections, and MEMCover, which integrates mutual exclusivity among genetic alterations with connectivity derived from PPI networks [27]. These methods however do not include additional layers of biological information directly into their model, such as RNA regulation or gene expression. Few approaches address the problem of integrating such additional data sources, among which TieDIE [37], which finds one large PPI subnetwork connecting DNA, RNA and regulatory signals. A separate class of data integration methods aiming to identify dysregulation in cancer focus on individual driver genes, without also connecting the drivers into modules, such as DriverNet [3], which combines mutations and gene expression, or DawnRank [25], which integrates mutations, gene expression and protein interactions.

Here, we describe ModulOmics, a method for the *de novo* identification of cancer driver modules based on the integration of PPI networks, mutual exclusivity of DNA alterations (SNVs and CNAs), and RNA-level co-regulation and co-expression, into a single probabilistic score (Figure 1). We identify modules that maximize this score by performing a two-step optimization procedure that combines Integer Linear Programming (ILP) with stochastic search. We apply ModulOmics on three large-scale TCGA datasets of breast cancer [5], glioblastoma (GBM) [6] and ovarian cancer [7], and show that it accurately identifies known cancer driver genes and pathways. Moreover, ModulOmics outperforms three state-of-the-art methods to detect cancer modules, namely the DNA-centric method TiMEx [12], the PPI-based method HotNet2 [29] and the DNA and PPI integration method MEMCover [27].

**Figure 1:**
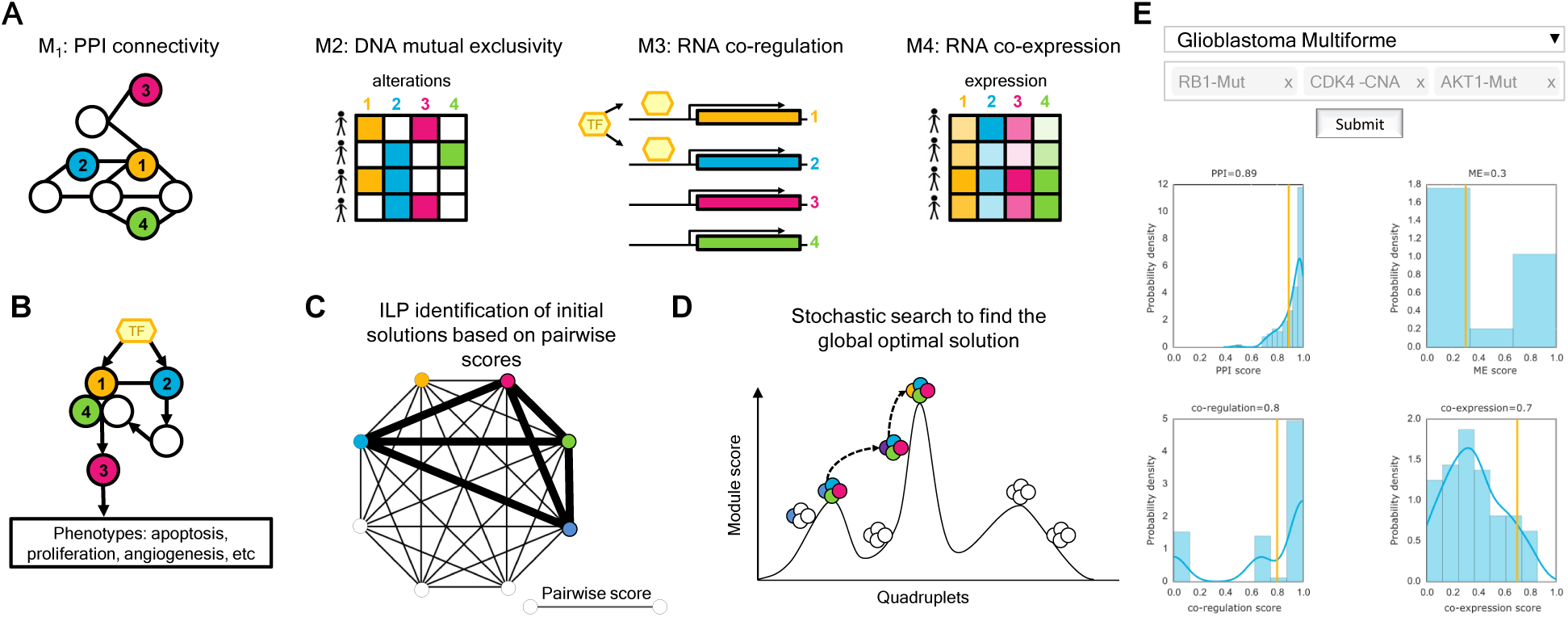
Overview of ModulOmics. **A)** Four different data sources, corresponding to four different models *M*_1_, …, *M*_4_ (see Methods), contribute to the computation of the ModulOmics score: PPI connectivity (protein level), mutual exclusivity (DNA level), transcriptional co-regulation (regulatory connections and RNA level) and co-expression (RNA level). The four colors correspond to four different genes; full squares in the matrix for model *M*_2_ encode the presence of alterations, while empty squares encode their absence. In *M*_3_, genes 1 and 2 are regulated by a common transcription factor. In *M*_4_, the different color intensities depict different expression intensities. **B)** Potential mechanism leading to a driver module exhibiting patterns of PPI connectivity, mutual exclusivity, co-regulation and co-expression. **C)** The ILP optimization identifies modules with highest sum of pairwise ModulOmics scores, computed as the average of the four scores corresponding to models *M*_1_, …, *M*_4_, further z-scored and normalized to [0,1]. **D)** The stochastic search optimization uses the modules identified by ILP, depicted in panel C, as seeds, and aims to improve their scores and identify the global optimal solution. The space of initial solutions is clustered and genes are exchanged between clusters in order to identify modules with high global scores. While the scores for models *M*_1_, …, *M*_4_ of the modules in panel C were approximated as the average pairwise scores, here they are computed explicitly for the entire module. **E)** The webserver tool computes the ModulOmics score of any chosen gene set, based on any of the TCGA datasets analyzed in this study. For each data source employed by ModulOmics, the tool plots the single omics scores of the top 50 modules, highlighting the score of the chosen gene set.

We further use ModulOmics to identify modules that characterize breast cancer subtypes. The highest scoring modules are enriched with cancer drivers, and reliably separate cancerous from normal tissues in an independent patient cohort [40, 51]. Moreover, the modules characterizing aggressive subtypes, such as Her2 and Basal, are further enriched with Gene Ontology (GO) terms related to cell proliferation. In the triple negative (TN) subtype, we identify functional connections among multiple down-regulated tumor suppressors, including *TP53*, *BRCA1*, *RB1* and *PTEN*. This pattern is also supported by reverse phase protein array (RPPA) data [5]. In Luminal A, high scoring modules containing *PTEN* suggest two potential functionalities of this protein: a canonical one as part of the PI3K pathway, and a non-canonical one as a regulator of cell proliferation.

ModulOmics is freely available in two forms, as an open-source R code for the identification of cancer driver modules from a cohort of cancer samples (https://github.com/danasilv/ModulOmics), and as a webserver for the evaluation of any set of genes of interest using the TCGA data processed in this study (http://anat.cs.tau.ac.il/ModulOmicsServer/).

## Results

ModulOmics identifies driver modules on the basis on DNA and RNA cancer patient data, integrated with PPI networks and known regulatory connections. Each candidate module is scored according to the degree of mutual exclusivity among DNA alterations in its members across the patient cohort, the correlation of the RNA expression of its members across the cohort, the probability that its gene members are connected in the PPI network, and the fraction of its members that are co-regulated by a common active transcription factor. As the number of candidate groups grows exponentially with maximal group size, ModulOmics uses a heuristic two-step optimization procedure to first find good initial solutions by linearly approximating the scoring function and then refining these solutions via stochastic search (see STAR Methods). Each module is assigned an empirical p-value by comparing its score to the scores of 1,000 random modules of altered genes of the same size. All top modules identified by ModulOmics were significant (corrected Bonferroni p-value *<* 0.05).

We applied ModulOmics on three large TCGA cancer datasets: breast cancer, GBM, and ovarian cancer. On these datasets, we compared ModulOmics to four simplified similar approaches, in which the score of a group is computed using only single omic data sources, namely PPI connectivity, mutual exclusivity, co-regulation or co-expression, as well as to three state-of-the-art methods for the identification of driver modules: MEMCover, HotNet2 and TiMEx. We additionally assessed the contribution of each of the single omic data sources to the ModulOmics score by only using subsets of three omics, and removing each single omic at a time.

We found that modules of sizes 3 and 4 generally have higher ModulOmics scores than pairs (Figure S1), and no single omic data source dominates the score for any given module size (Tables S1-S3). Sensitivity analyses showed that the performance of ModulOmics is robust under different parameter choices (Supplementary Information, Figures S2-S3 and Tables S4-S5).

### Driver modules are enriched with cancer drivers

To assess the performance of ModulOmics, we calculated the enrichment of the highest scoring driver modules with known driver genes (positive controls) and known non-driver genes (negative controls). To this end, we used the gene lists introduced in [23], complied from different sources: the Network of Cancer Genes (NCG) [1], Cancer Gene Census version 73 (CGC) [17], the Atlas of Genetics and Cytogenetics in Oncology and Hematology (AGO) [26], UniprotKB [53], DISEASES [39] and MSigDB [47] (Supplementary Information). The enrichment was calculated as the fraction of gene members in each module that were also part of each control list, averaged across the top modules considered. The top 10 modules inferred by ModulOmics generally outperformed the top 10 modules identified with the four single omic approaches and with MEMCover, HotNet2 and TiMEx across the seven positive and three negative control lists tested (Figure 2A). Specifically, ModulOmics achieved an enrichment score of close to 1 across all three cancer types in the three largest positive control lists: the manually curated resource *NCG5*, the positive AGO list (*PosAGO*), and the Union All list (*PosUnionAll*), consisting of between 1,429 and 2,144 known drivers. Importantly, the modules inferred by ModulOmics scored close to 0 in all three negative control list assessed, namely the complete negative AGO list (*NegAgoFull*), the curated negative AGO list (*NegAGOClean*), and the negative list introduced in [15] (*NegDavoli*), consisting of between 3,272 and 9,457 known non-driver genes.

**Figure 2:**
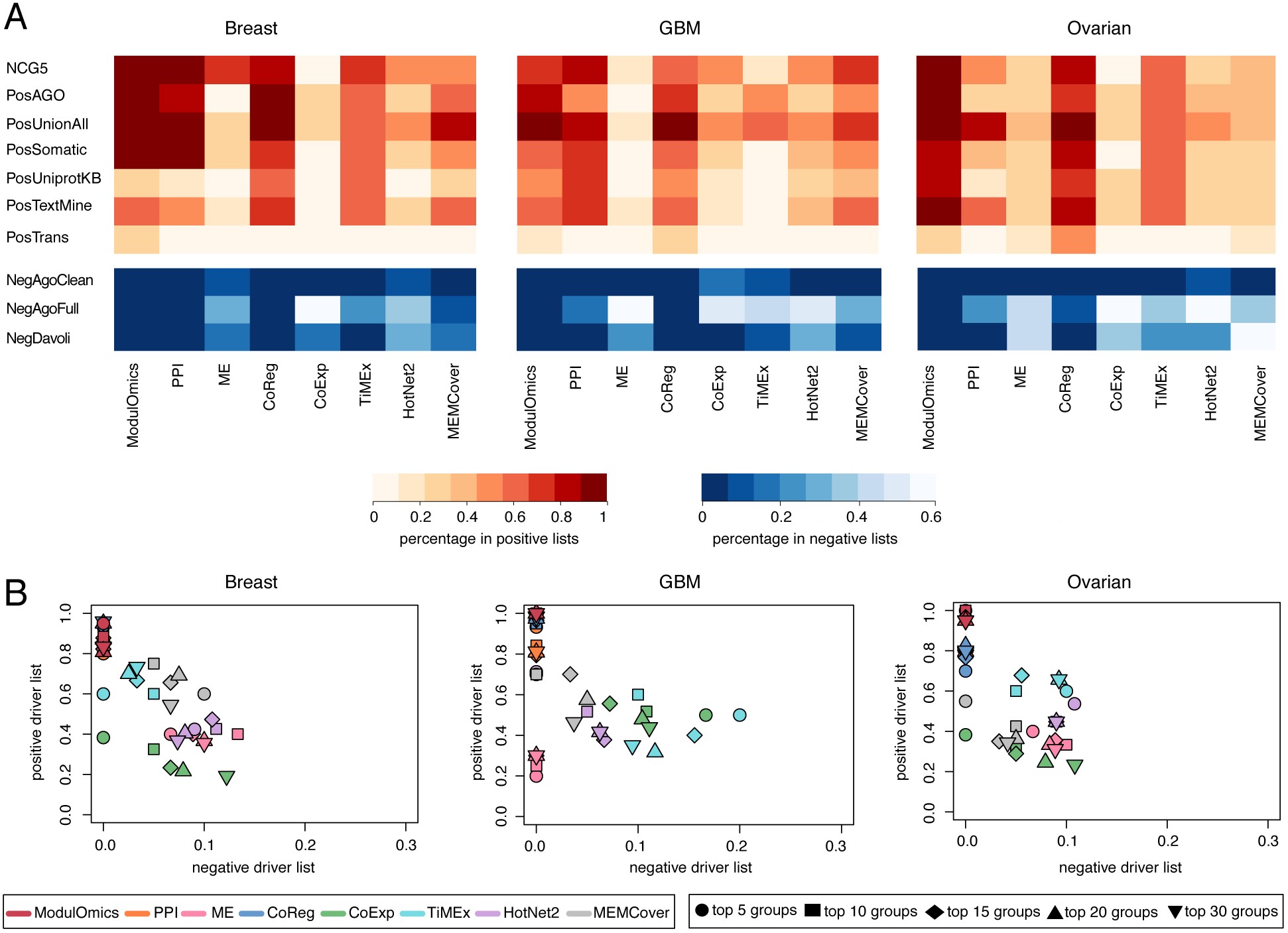
The driver modules inferred by ModulOmics are enriched with cancer driver genes. **A)** The average driver enrichment (red heatmaps) and non-driver enrichment (blue heatmaps) across the top 10 scoring modules inferred by each method in the three cancer types studied. The enrichment was calculated as the fraction of gene members in each module that are also part of each control list, averaged across the top 10 modules. The modules were ranked by their score, regardless of their sizes (the inferred groups consisted of two, three or four gene members). *ME* stands for mutual exclusivity, *CoReg* for co-regulation, and *CoExp* for co-expression single omic scores. *NCG5*, *PosAGO*, *PosUnionAll*, *PosSomatic*, *PosUniprotKB*, *PosTextMine*, *PosTrans* are the positive control lists, while *NegAgoClean*, *NegAgoFull* and *NegDavoli* are the negative control ones (see STAR Methods). **B)** Detailed driver and non-driver enrichment scores for the positive driver list *PosUnionAll* and the negative driver list *NegAGOClean* for the seven methods assessed, across the three cancer types, for the top scoring 5, 10, 15, 20 and 30 modules. Table S6 shows the scores for 2B.

In addition, ModulOmics also outperformed the other methods when evaluating the highest scoring 5, 10, 15, 20, or 30 modules of any size (Figure 2B), or when separately evaluating modules of fixed sizes (Figure S4). Among the competing methods, PPI-based and co-regulation-based scorings exhibited good performances, MEMCover performed well only in the case of certain group sizes, while co-expression, mutual exclusivity, HotNet2 and TiMEx generally performed poorly on both positive and negative control metrics. We further evaluated the contribution of each single omic data source to the driver genes enrichment by computing reduced versions of the ModulOmics score, each time with a single omic removed. We found that integrating all four omics data sources improves the enrichment, as compared to using subsets of three omics, in 90% of the evaluated cases (92% of the positive control lists and 86% of the negative lists). Nevertheless, the performance of ModulOmics remained fairly robust when using only three omics sources, suggesting that the method can also be applied in cases when one data source is missing (Figure S5).

One of the features of ModulOmics is that each gene can participate in multiple pathways, hence the reported modules often overlap (Figure S6). Biologically, this feature is justified by the fact that the known driver genes are likely network hubs, expected to be functionally connected to multiple other less-known driver genes into different modules. In order to assess the performance of ModulOmics also in the absence of overlap among groups, we repeated the driver evaluation while considering the first 20 unique appearances of genes in the top modules. Consistent with the previous results, ModulOmics outperformed the three other competitive methods tested (Figure S7).

### Driver modules are functionally coherent

An additional metric for evaluating the relevance of the inferred modules concerns their functional coherence, which we assessed via their enrichment with curated pathways from KEGG [33] (Supplementary Information). ModulOmics identified key cancer-related pathways, such as *pathways in cancer*, across all three cancer types (Figure 3A), without showing preferential enrichment for particular module sizes (Figure S8). In contrast, HotNet2 identified this pathway only in the GBM and ovarian cancer datasets, and MEMCover and TiMEx did not identify it at all. Additional highly enriched pathways included *apoptosis*, *cell cycle*, *TP53 signaling*, *mTOR signaling*, and the angiogenesis-related *VEGF pathway*. Interestingly, the set of enriched pathways also included pathways characterizing other cancers types, indicating shared mechanisms among malignancies.

**Figure 3:**
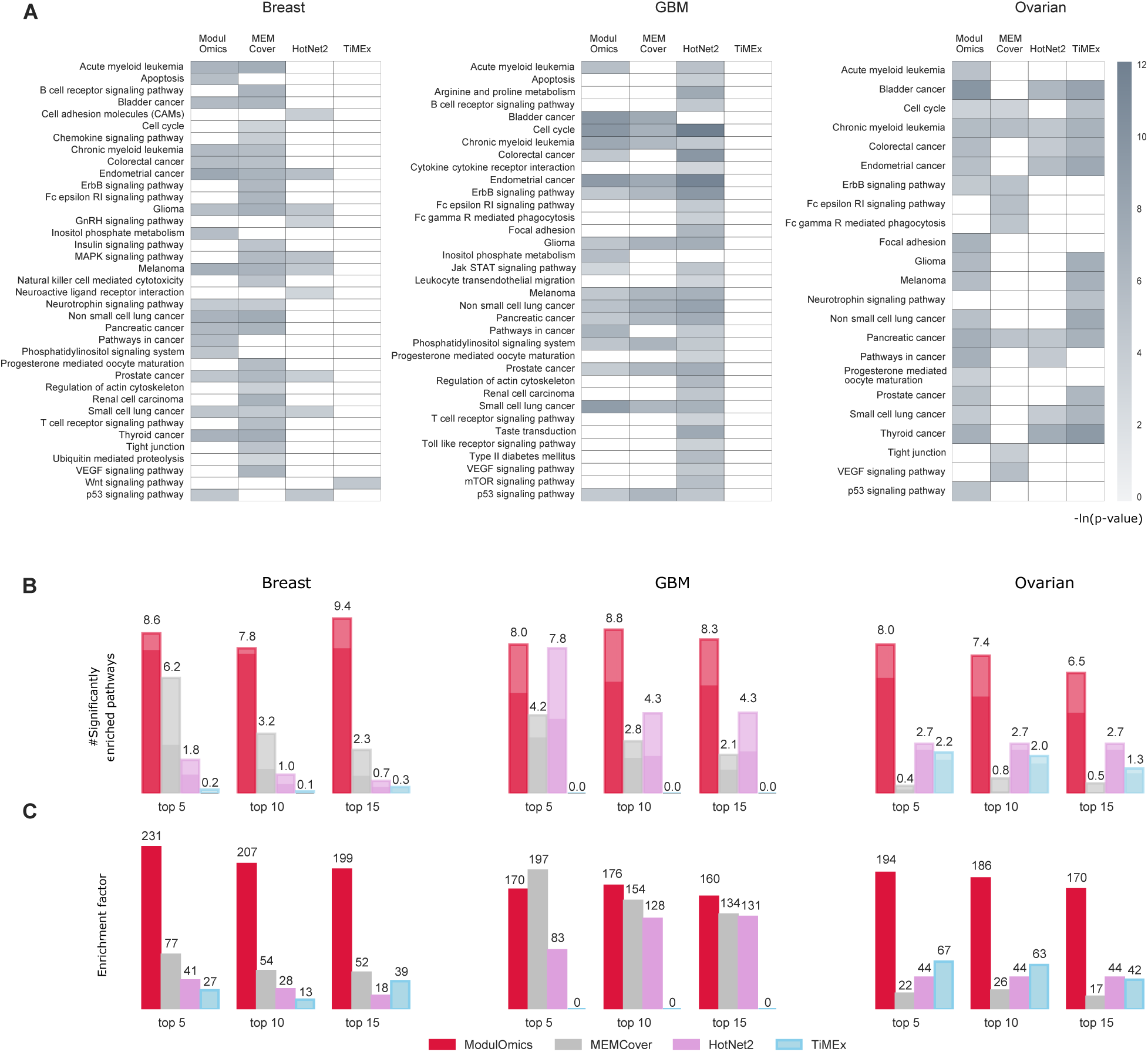
The driver modules inferred by ModulOmics are enriched with cancer driver pathways. **A)** Mean hyper-geometric p-value of the KEGG pathways significantly enriched in the top 10 modules identified by ModulOmics (red), HotNet2 (purple) and TiMEx (light blue). **B)** Average number of KEGG pathways significantly enriched in the top modules, indicated above the bars. The opaque bars indicate cancer-related pathways only. **C)** Average enrichment factors for top modules. The numbers displayed in panels B and C are normalized per module. Enrichment p-values and factors were computed with Expander [52].

To quantify the pathway enrichment performance of ModulOmics, MEMCover, TiMEx and HotNet2, we counted the number of pathways significantly enriched (Bonferroni-corrected p-value ≤0.05) in each of the top 5, 10 and 15 highest scoring modules (Figure 3B), and computed their average enrichment factor (Figure 3C). Enrichment p-values and factors were computed with Expander [52] (Supplementary Information). Overall, ModulOmics identified modules enriched with more general pathways and cancer-related pathways than the three competing methods, and was the only method for which all highest scoring 10 modules were enriched with at least one pathway. A high percentage of the genes identified by ModulOmics participated in known KEGG pathways, reaching an average of 78% across all three cancer types, compared to 43%, as identified by MEMCover, 39% by HotNet2, and 13% by TiMEx (Table S7). In contrast, less than 5 out of the 1,000 random modules generated for computing module significance were significantly enriched with any known pathway. Finally, we tested the contribution of each omic to pathway enrichment and found that using all four data sources improves the identification of functionally coherent modules in 92% of the tested cases (Figure S9).

### Driver modules in breast cancer subtypes recapitulate known mechanisms and suggest novel functionalities

Next, we applied ModulOmics on molecularly defined subtypes of breast cancer, classified using the mRNA PAM50 classification [36] into Basal (125 patients), Her2 (61), Luminal A (364) and Luminal B (174) (Table S8 and Figure S10). Across all subtypes, the genes in the top 20 modules (Figure 4A) were highly enriched with cancer drivers (66% were part of the NCG5 positive control list and 70% were part of the UnionAll positive list, while only 4% were part of the AGOClean negative control list) and KEGG pathways (44 enriched pathways, 24 of which were directly related to cancer, average p-value 0.0063). The top drivers identified by ModulOmics included *TP53*, *AKT1*, *mTOR* and *PTEN*, as well as subtype-signature genes such as *BRCA1* and *BRCA2* for Basal [49, 50], *CDH1* for Luminal A and B [24], *MAP3K1* for Luminal B [5] and *EGFR* for Her2 [32]. An alternative strategy to ModulOmics for identifying relevant drivers would have been selecting genes with highest SNV or CNA alteration frequencies [54]. However, in that case, a substantial portion of the enriched gene list identified by ModulOmics would have been overlooked, as 34% fall below the SNV median frequency per gene and 40% fall below the CNA median frequency per gene (Figure 4A). Therefore, integrative approaches such as ModulOmics are essential to this end.

**Figure 4:**
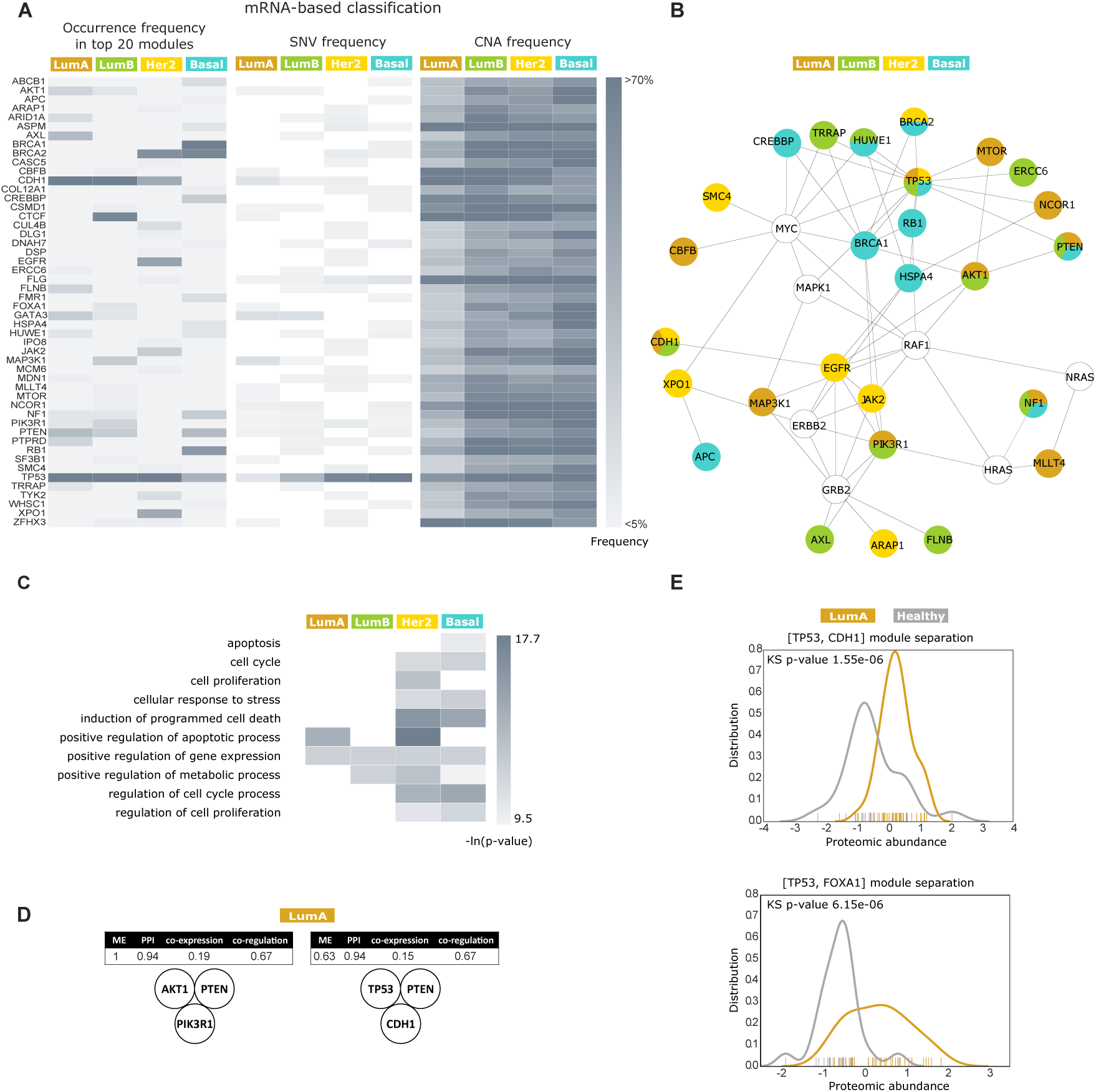
Modules inferred in mRNA-classified breast cancer subtypes reflect various levels of subtype aggressiveness and separate cancerous form healthy tissues. **A)** For each mRNA-based subtype and for the pooled set of genes in the top 20 modules, we computed their occurrence frequency in the top 20 modules, as well as their SNV and CNA alteration frequencies across the patient cohort. These genes are enriched with known cancer drivers and pathways, and could not have been identified if relying on SNV and CNA alteration frequencies alone. White corresponds to absent genes (0% frequency). **B)** Detailed PPI network view of the subset of genes in panel A that are either known drivers, or part of KEGG pathways. The displayed physical protein interactions underline cancer-related functional associations, such as the role of the PIK3A pathway in Luminal A tumors. **C)** Selected list of significantly enriched GO pathways across the top 20 modules (Figure S11 displays the full list), reflecting the aggressiveness of the Basal and Her2 subtypes, compared to Luminal A and Luminal B. Enrichment hyper-geometric p-values were computed with Expander [52]. White corresponds to absent pathways. **D)** Module scores for top Luminal A modules suggesting two different biological roles of the tumor suppressor *PTEN*. **E)** The highest ranking Luminal A module in an independent mass spectrometry proteomics dataset separates cancerous from healthy patient tissues. *TP53* loss is measured by its downstream regulated protein CDC2, *CDH1* loss is measured by its downstream regulated protein CTNNB1, and *FOXA1* gain is measured directly.

A detailed PPI network view of the genes identified by ModulOmics showed that *TP53* is a key player in tumor progression for all subtypes, while subtype-specific key players included *EGFR* for Her2 and *BRCA1* for Basal (Figure 4B). The network view highlighted the higher rate of PPI-connected established tumor suppressors in the Basal subtype, as compared to Luminal A, matching the aggressive nature of these tumors. In addition, Luminal A modules were characterized by a higher occurrence of PI3K pathway members, such as *PIK3R1*, *AKT1*, *mTOR* and *PTEN*, as previously observed [5]. The top modules identified by ModulOmics were further highly enriched with functional relations, highlighting different GO annotations for each subtype (Figure 4C). These results captured the increased pathway activity of key pathways required for tumor progression, such as apoptosis, cell cycle process or cell proliferation, as well as the known aggressiveness of Basal and Her2 tumors, reflected in their higher pathway enrichment.

The highest ranking module for both Luminal A and Luminal B was *TP53* and *CDH1*, two known functionally associated drivers in the Luminal subtypes [5, 4]. The top Her2 modules were characterized by the recurrent appearance of the nuclear export gene *XPO1* together with *TP53*, which is one of its known targets [18, 9]. Interestingly, the mechanism of TP53 nuclear export by *XPO1* is well characterized [16], yet no pivotal role was suggested specifically in Her2 breast cancer. The highest ranking module of the Basal subtype consisted of *RB1*, *BRCA1*, *NF1*, and *CREBBP*. *CREBBP* is a *BRCA1* activator [35] and, since both *BRCA1* and *CREBBP* are involved in DNA repair, this module potentially reflects the altered DNA damage repair mechanism specific to Basal tumors [34].

One of the frequently occurring genes in the top Luminal A modules was the tumor suppressor *PTEN*, occurring both in modules reflecting its canonical PI3K pathway role, and in modules suggesting a non-canonical role (Figure 4D). The canonical *PTEN* module also included *PIK3R1* and *AKT*, thus supporting the known mutual exclusivity pattern of mutations within the PI3K pathway [42]. The module suggesting the non-canonical role of *PTEN* also included *CDH1* and *TP53*, supporting the hypothesis that *PTEN* regulates cell proliferation by increasing the binding of *CDH1* to APC\C, a complex known for its tumor-suppressive function, and by increasing *TP53* acetylation following DNA damage [44]. Indeed, according to the TRRUST database [21], *PTEN* and *CDH1* are co-regulated by two common transcription factors, namely *STAT3* and *NFKB1*.

In order to further explore the clinical relevance of the highest scoring driver modules, we examined how well they can distinguish healthy tissues from cancerous ones in an independent omic data source. To this end, we used a recently published proteomics dataset consisting of 62 samples of Luminal A and healthy tissues [40, 51], and focused on the two highest scoring Luminal A modules: *TP53* and *CDH1*, and *FOXA1* and *TP53*, respectively. These top two modules significantly separated the Luminal A cancerous tissues from the healthy ones (p-value 1.6*e*^−06^ and p-value 6.2*e*^−06^ respectively, KolmogorovSmirnov (KS) test, Figure 4E). For comparison, neither *GATA3* or *PIK3CA*, the most frequently mutated genes in Luminal A, nor *TP53*, the most frequently mutated gene in breast cancer, were able to significantly separate the two types of tissue (p-value 0.065, p-value 0.054, and p-value 0.69, respectively, KS test). Similarly, random modules of the same size did not significantly separate the tissues (p-value 0.14, averaged over 1,000 random modules generated by sampling subsets of proteins from the proteomics dataset, KS test).

An alternative way to study breast cancer progression is by stratifying patients according to immunohistochemistry results assessing the HER2, ER and PR receptors (Supplementary Information). To this end, we separated our patient cohort into the following subtypes: TN (116 patients), Her2-enriched (30), Luminal A (477) and Luminal B (88), and used ModulOmics to infer modules for each subtype (Table S9 and Figure S12). Similarly to the mRNA-based classification, the genes part of the highest scoring 20 modules were enriched with cancer drivers (67% were part of the NCG5 positive control list and 59% were part of the UnionAll positive list, while only 2% were part of the AGOClean negative control list) and with known cancer pathways (46 enriched pathways, 25 of which were directly related to cancer, average p-value 0.01, Figure S13A). Across subtypes, the highest scoring modules highlighted a unique alteration pattern for the tumor suppressor *TP53*. In Luminal A, Luminal B and Her2-enriched, *TP53* was mutually exclusive with other tumor suppressors, such as *PTEN* and *BRCA1* in Luminal B, or *BRCA2* in Luminal A, which led to ModulOmics inferring these groups as high scoring modules. However, in TN, *TP53* was mutually exclusive with *BRCA2*, but not with other key TN drivers, such as *BRCA1*, *PTEN* or *RB1*, as both the pairwise and the group mutual exclusivity scores of *TP53* and these three drivers were 0 (Figure S13B). This suggests a TN-specific concerted down-regulation of multiple tumor suppressors, namely *TP53*, *BRCA1*, *RB1* and *PTEN*, potentially contributing to the bad prognosis of this subtype. Taken together, these results imply that the level of mutual exclusivity in tumor suppressors might reflect the aggressiveness of the tumor subtype [38, 45, 5, 14].

Finally, we used an independent omic data source (RPPA) to further evaluate the functional connectivity among the tumor suppressors *PTEN*, *BRCA1*, *RB1*, and *TP53*. In general, evaluating protein measurements limits automatic and exhaustive analyses, since loss of function can lead to missing data, requiring the identification of downstream regulated proteins that can serve as surrogates. PTEN was found to be downregulated, while phosphorylated AKT, which is suppressed by PTEN, was upregulated. *BRCA1* loss was accounted for by its downstream regulated protein CYCLIN B1, which was highly expressed. RB1 showed an overall low expression in most samples, while the RB1-related CYCLIN-D1 was lowly expressed mostly in TN tumors (Figure S13C). *TP53* loss was accounted for by its target CDK1, which was also highly expressed. *CDK1*, *CYCLIN B1* and *AKT*, the tumor promoters regulated by *TP53*, *BRCA1* and *PTEN*, were significantly upregulated in TN tissues compared to the other subtypes (p-value 1.2*e*^−16^, KS test), while the tumor suppressors *PTEN*, *RB1*, and *BRCA2* were significantly downregulated (p-value 2.9*e*^−09^, KS test, Figure S13D). These results suggest that these two groups of genes can be used to separate TN from the other subtypes.

## Discussion

ModulOmics is a novel method to *de novo* identify molecular cancer driver pathways, based on the integration of connectivity within protein-protein interaction networks, mutual exclusivity among SNV or CNA alterations, transcriptional co-regulation, and RNA co-expression, into a single probabilistic score. ModulOmics uses an efficient two-step optimization procedure to first find good initial solutions using linear approximation, and then refine these solutions with stochastic search. We demonstrate the performance of ModulOmics in identifying modules enriched with known cancer driver genes and pathways in three large-scale multi-omics TCGA datasets: breast cancer, GBM, and ovarian cancer. We further investigate breast cancer subtypes and find that some of the highest scoring modules are known to be involved in cancer-related molecular mechanisms, while others suggest novel functionalities. We evaluate these results using an independent patient cohort and independent proteomic and phosphoproteomic datasets. In addition, we show that the top modules inferred by ModulOmics can be used to reliably separate cancerous from normal tissues in Luminal A samples, as well as to distinguish TN samples from the other subtypes.

ModulOmics is a freely available open-source software. The framework is designed to be flexible, such that any of the four sources of evidence employed here can be excluded or replaced with new sources of evidence. In addition, since ModulOmics integrates independent sources of information, newly added data can also originate from different patient cohorts, such that single omic datasets from other cancer studies can be readily integrated. Moreover, the webserver implementation of ModulOmics can be used to evaluate the ModulOmics score of any user-defined gene set, on the basis of any of the TCGA datasets analyzed here. This application can be very useful in situations when gene sets were deduced from separate biological or computational analyses.

Some of the modules identified by ModulOmics may merit further experimental investigation. For example, on the basis of the highest ranking Basal module (*RB1*, *BRCA1*, *NF1*, and *CREBBP*), we propose that further validation experiments could evaluate the clinical implications of using the *CREBBP* inhibitor in *BRCA1* patients, similarly to *PARP1*, another DNA repair agent successfully used in treatment [28]. Based on the recurrent joint occurrence of *XPO1* and *TP53* in top Her2 modules, we propose to further evaluate the role of the export mechanism of *XPO1* leading to *TP53* depletion in the nucleus, thus decreasing its tumor suppression capability. The role of *XPO1* in tumor progression was previously investigated in a preclinical context of TN treatment [10, 31]. Here, we suggest it may also play a role in the Her2 subtype. Additionally, in the future, once high-throughput single-cell profiling of tumors becomes routinely performed in the clinic, ModulOmics can be used to identify functional connections derived from multiple tumor cells of single cancer patients, rather than a patient cohort. In this way, the design of personalized cancer therapies based on the tumor heterogeneity of each patient can be facilitated.

A unique feature of ModulOmics is that its scoring function not only uses different types of data, but also integrates different types of statistical tests distinctly designed for each data type (such as the mutual exclusivity test for mutational data, or the proximity-based PPI score). In contrast, previous approaches generally apply the same methodological framework to all data types [37]. Integrating different statistical tests is empowered by online normalizing each score throughout the search, as well as by optimizing the different scores simultaneously, rather than sequentially. In addition, the weights of the different scores can be optimized from the data, with the aim of identifying groups representative of a functional phenotype of interest. To exemplify this, we inferred optimal omics weights by training a classifier to identify groups enriched with the Cell Cycle GO term from the breast cancer top modules reported by ModulOmics. The classifier assigned the weight 0.61 to the ME score, 3.61 to the PPI score, 1.83 to the co-regulation score and 1.93 to the co-expression score. All the four data sources were hence found to contribute to the classification of modules associated with the GO term, with the scores reflecting physical interactions carrying more weight, as would be expected in the case of phenotypes related to biological processes.

ModulOmics can be extended beyond finding modules in cancer. For example, the PPI, co-regulation and co-expression scores can be used to detect protein complexes. As a proof-of-concept, we collected 1,112 known biological complexes from CORUM [41] (see details in the Supplementary Information) of sizes 2, 3 and 4, as well as equal number of random protein groups and calculate their ModulOmics scores. We trained a classifier to distinguish known complexes from random groups based on the three scores, reaching an AUPR of 0.84 for all module sizes (Figure S14) and of up to 0.9 for modules of size 4. These results indicate that the ModulOmics scores are informative in identifying new biological connections outside the scope of cancer, making the tool broadly applicable.

## STAR Methods

### Model

Given a set *G* = {*G*_1_, …, *G*_*n*_} of genes and a collection *M* = {*M*_1_, … *M*_*m*_} of models for different data types, we are interested in computing *S*_*G*_, the ModulOmics probabilistic score of the set *G*, reflecting how likely are the genes in *G* to be functionally connected. *S*_*G*_ is computed as the mean of *m* probabilistic scores *P* (*G* | *M*_*k*_). Each of these *m* scores represents how strongly functionally connected the genes in *G* are, under different models:

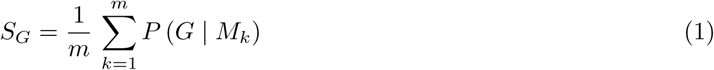

The models we consider here are: connectivity among protein-protein interactions (*M*_1_), mutual exclusivity among point mutations or copy number alterations (*M*_2_), transcriptional co-regulation (*M*_3_), and gene coexpression (*M*_4_).

### PPI Connectivity

Model *M*_1_ assesses the functional connectivity of the set *G* at the protein level, by computing the probability of *G* being connected in the PPI network. Starting with a fully-connected literature-based PPI network (HIPPIE) and its associated interaction probabilities, we define, for each pair of genes (*G*_*i*_, *G*_*j*_), con (*G*_*i*_, *G*_*j*_) as the probability of the most likely path connecting *G*_*i*_ and *G*_*j*_, *i.e.*, the product of the probabilities of the path’s edges. The computation of con(*G*_*i*_, *G*_*j*_) for all *G*_*i*_, *G*_*j*_ ∈ *G* yields a complete graph on *G*, denoted 𝒢(*G*). If we denote the edge set corresponding to any graph *H* by *E*(*H*), then the connectivity of the set *G* is defined as the sum of the probabilities over all connected subgraphs spanning *G*, as follows:

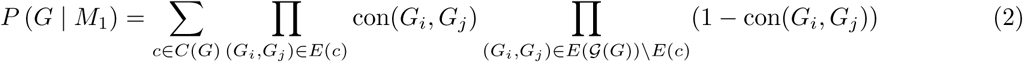

where *C*(*G*) is the collection of connected subgraphs spanning 𝒢(*G*).

### Mutual exclusivity

Model *M*_2_ estimates the degree with which DNA alterations support the functional connectivity of the genes in *G*. Following the mutual exclusivity framework defined in the context of waiting times to alteration introduced in TiMEx [12] and pathTiMEx [13], *P* (*G* | *M*_2_) is computed as the degree of mutual exclusivity of the set *G*, as follows:

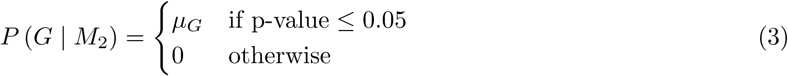

where both *µ*_*G*_ and p-value are reported by TiMEx. The TiMEx probabilistic graphical model estimates *µ*_*G*_, which is the mutual exclusivity intensity of the group *G*, via a nested likelihood ratio test between an independence model and an alternative, mutual exclusivity model. The independence model assumes that the genes evolve independently during disease progression, whereas the mutual exclusivity model assumes that only the gene with the shortest waiting time in a functionally connected group of genes will fixate. The parameter *µ*_*G*_ represents the probability that a group of genes is perfectly mutually exclusive, *i.e.*, that no two genes in *G* share alterations in the same patient. Therefore, *µ*_*G*_ = 1 corresponds to perfect mutual exclusivity, and *µ*_*G*_ = 0 corresponds to independence. The p-value in Equation 3 is the probability of observing a given alteration pattern of the set *G* under the null hypothesis of independence, as described in [12] and [13].

### Co-regulation

Model *M*_3_ assesses the functional connectivity of the genes in *G* on the basis of their transcriptional regulation. The co-regulation score *P* (*G* | *M*_3_) is defined as the fraction of genes in *G* which are co-regulated by at least one common active transcription factor,

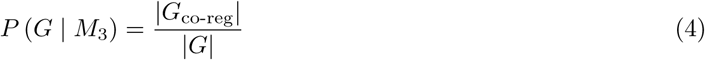

where *G*_co-reg_ ⊆ *G* is the maximal set in which all genes are regulated by at least one common active transcription factor. A transcription factor is considered active if it is differentially expressed (z-score of fold change is either *>* 1 or *<* −1) in at least 25% of samples. Alternatively, other operators such as the average could be used, however choosing the maximal set reflects the co-regulation of the entire group, rather than particular subgroups.

### Co-expression

Model *M*_4_ evaluates the functional connectivity of the genes in *G* based on their transcriptional profiles. Let a gene be defined as expressed if its expression averaged across all samples is above the *k*^th^ *q*-quantile, and let *G*_exp_ ⊂ *G* be the set of all expressed genes. Then, the co-expression score of *G* is defined as the mean among all pairwise Spearman correlations of the expression profiles of the genes in *G*_exp_, and 0 corresponding to the remaining pairs, in which at least one of the genes is not expressed,

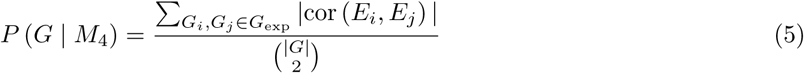

where *E*_*i*_ is the continuous expression level of gene *G*_*i*_ across all samples, and cor(*E*_*i*_, *E*_*j*_) is the Spearman correlation among the expression profiles of *G*_*i*_ and *G*_*j*_. For this application, we chose *k* = 2 and *q* = 4, *i.e.*, the 2^nd^ quartile. The choice of Spearman correlation is justified by not necessarily assuming a linear relation between expression profiles. Missing expression data can be handled by assigning the respective genes null expression profiles, leading to their consideration as unexpressed genes.

## Optimization procedure

Given a large cancer dataset, identifying groups of functionally connected genes is challenging, as the number of candidate groups increases exponentially with maximal group size. Therefore, we employ a two-step procedure to optimize the global ModulOmics score in equation 1. First, in order to identify a large set of good initial solutions, we formulate the optimization problem as an ILP, and optimize a linear approximation of the global ModulOmics score. Second, we perform a stochastic search starting from these initial solutions and using the global score.

### ILP

The first step of our optimization procedure linearly approximates the exact scores of the set *G* under each of the four models *M*_*k*_, by decomposing them into pairwise scores. For each model *M*_*k*_, the score of each pair of genes (*G*_*i*_, *G*_*j*_) is denoted by 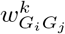 and equals to *P* ((*G*_*i*_, *G*_*j*_) | *M*_*k*_), further z-scored and normalized to [0,1]. The goal of the optimization routine is to identify candidate subsets *G* with high total scores *w*_*G*_, computed as:

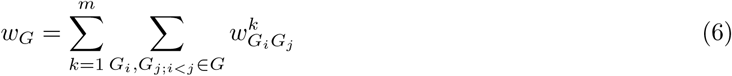

The ILP retrieves sets *G* of fixed size *K* with maximal *w*_*G*_ score. Thus, *G* is the maximal weight subgraph of size *K* in a weighted complete graph with vertices *V*, corresponding to a large set of genes, and edges 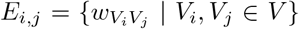. The ILP consists of the following set of binary vertex variables *V*_(*i*)_ denoting the inclusion of vertex *V*_*i*_ in a set *G*, and edge variables *E*_(*i,j*)_, denoting the inclusion of edge *E*_*i,j*_ in *G*:

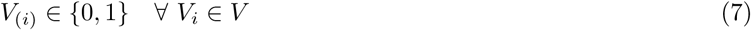

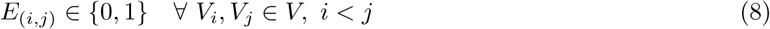

and the objective function:

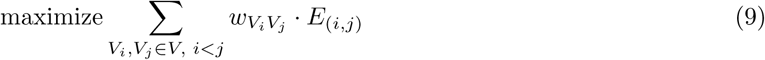

under the constraints:

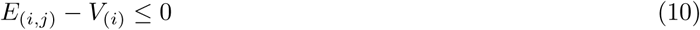

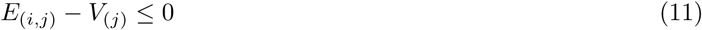

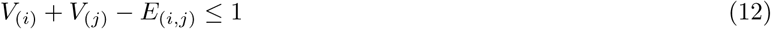

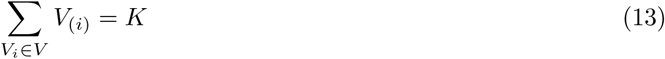

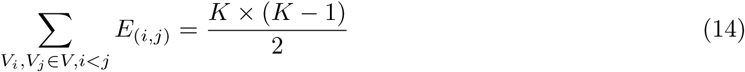

∀ *V*_*i*_, *V*_*j*_ ∈ *V*, *i* < *j*. Constraints 10, 11, and 12 ensure that the retrieved set is a clique, and constraints 13 and 14 ensure that the clique is of size *K*. Let us note that identical solutions would be retrieved by discarding either constraint 13 or 14, yet we include both for efficiency considerations. With each candidate set *G* found, we add constraint 15 to prevent the ILP to choose the entire set *G* again:

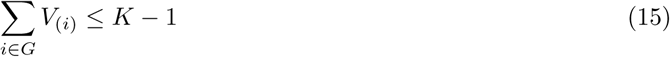

### Stochastic search

We use 200 high-ranking modules identified by the ILP as seeds for a stochastic search that expands the search space and optimizes directly the exact score of the modules, rather than their pairwise approximations. The stochastic search uses the seed modules as starting points and aims to find the modules with global optimal score by offering possible exchanges of module members. The seed modules are clustered into 10 clusters using *k*-means, and a search cycle starts independently from each cluster, in order to increase the chances of finding modules with global optimal scores. Each of these 10 cycles iterates among the modules in its cluster and tries to improve each one by suggesting 20 possible exchanges of a random module member with another random gene. If the score improves, then the exchange is accepted and the module is updated accordingly. Each cycle reports its 5 highest scoring modules. The modules reported by all 10 cycles are finally aggregated and re-ranked. Sensitivity analyses show that the performance of ModulOmics is robust under different parameter choices (Supplementary Information). Each run of the ILP followed by the stochastic search yields optimal modules of fixed size *K*. To retrieve the top modules in a range of sizes we run the tool with *K* ranges from 2 to 4, aggregate the results and retrieve the top modules regardless of their size.

