## Supplementary Materials for "ModulOmics: Integrating Multi-Omics Data to Identify Cancer Driver Modules"

### 1 Data processing

#### 1.1 Data acquisition

ModulOmics identifies driver modules on the basis on DNA and RNA cancer patient data, integrated with PPI networks and known regulatory connections. We analyzed patient data from the TCGA project [2, 3, 4], downloaded from cBio portal [5]. We integrated the patient data with the PPI network Hippie [18] containing 238,165 physical interactions, and with the regulatory connections from the database Transcriptional Regulatory Relationships Unraveled by Sentence-based Text Mining (TRRUST) [9] containing 8,908 Transcription Factor (TF)-target regulatory relationships of 821 human TFs.

To further evaluate functional connections in the triple negative (TN) breast cancer subtype, we used the reverse phase protein array (RPPA) data published by the TCGA. In addition, we evaluated the power of the highest ranking modules in distinguishing healthy tissues from cancerous ones using an independent mass-spectrometry dataset containing over 62 samples of Luminal A and healthy tissue [16, 20]. Proteins were quantified using the method Super-SILAC [8].

#### 1.2 Criteria for receptor-based classification

Subtyping of breast cancer in clinical practice is mostly done by immunohistochemistry (IHC), based on the expression of estrogen receptor (ER), progesterone receptor (PR), and human epidermal growth factor receptor 2 (Her2). In this study, we classified the samples based on the information available in TCGA. The ER positive group was stratified based on Her2 expression, into Luminal B subtype for cases with positive receptor expression, and Luminal A for cases with negative expression. The ER negative group was segregated based on Her2 expression as Her2-amplified subtype in case of positive expression and TN subtype in case of negative expression.

### 2 Comparison with other tools

#### 2.1 HotNet2

HotNet2 [13] identifies subnetworks of a PPI that contain genes with significant numbers of mutations. HotNet2 uses a localized heat diffusion process to combine the mutational genomic data with the PPI data. To calculate statistical significance, HotNet2 uses a permutation test in which the gene scores are permuted among the genes, followed by a two-stage statistical test: first, a p-value for the number of subnetworks in the list is computed, and second, the estimate of the false discovery rate of the list of subnetworks is obtained. As recommended by the authors, we used SNVs and CNAs as the prior set, and assigned the initial prior score of each genetic alteration to be its alteration frequency in the data. We applied HotNet2 on the same PPI network we used with ModulOmics. To assign a p-value, we used 100 permuted networks as background. To calculate a hyper-geometric score for pathway enrichment with Expander [21], we used modules of up to size 7, since larger modules are more likely to be unspecific from a mechanistic perspective.

#### 2.2 TiMEx

We ran TiMEx [6] with default parameters on the same binary dataset used as input for ModulOmics, consisting of binary SNVs and CNAs alterations. We considered as significant all resulting mutually exclusive groups with Bonferroni-corrected p-value $< 0.1$. Even though TiMEx and the mutual exclusivity score of ModulOmics are based on the same probabilistic model, the search strategy is different for the two methods. Therefore, TiMEx and the simplified omic approach of mutual exclusivity are expected to identify different modules in the data.

#### 2.3 TieDIE

We compared our results with TieDIE [14], a method to detect one single subnetwork by incorporating mutation frequency derived from DNA data, differential expression derived from RNA data, regulatory network information, and a PPI network. TieDIE amplifies the signal from different inputs throughout the PPI network using network propagation and extracts the subnetwork casted by the amplified signals. ModulOmics differs from TieDIE in its goal, as TieDIE detects a single subnetwork, whereas ModulOmics identifies multiple modules.

We ran TieDIE on the three cancer cohorts used in this study: GBM, breast and ovarian cancers, using the same PPI and regulatory networks as for ModulOmics. ModulOmics modules were consistently enriched with over 80% of driver genes and under 5% of non-driver genes, while the subnetwork inferred by TieDIE, consisting of more than 300 genes, contained less than 30% known driver genes and more than 10% known non-driver genes. In addition, the KEGG pathway enrichment scores of the modules identified by ModulOmics ranged from 160 to 231, while TieDIE scores ranged from 5 to 76, across the three cancer types.

#### 2.4 MEMCover

We ran MEMCover [12] on each of the three cancer types, with default parameters. We used the same PPI as for ModulOmics, with an edge weight threshold of 0.4. The resulting modules were separated by size and further ranked by their average coverage in each cohort.

### 3 Extended Results

#### 3.1 Module size statistics

ModulOmics identifies modules of fixed sizes (in this application, we used 2, 3, and 4), which are further merged and ranked according to their scores. Figure S1 shows the distribution of module sizes across the top 50 modules, while tables S1- S3 show the average single-omic and multi-omics scores for each module size.

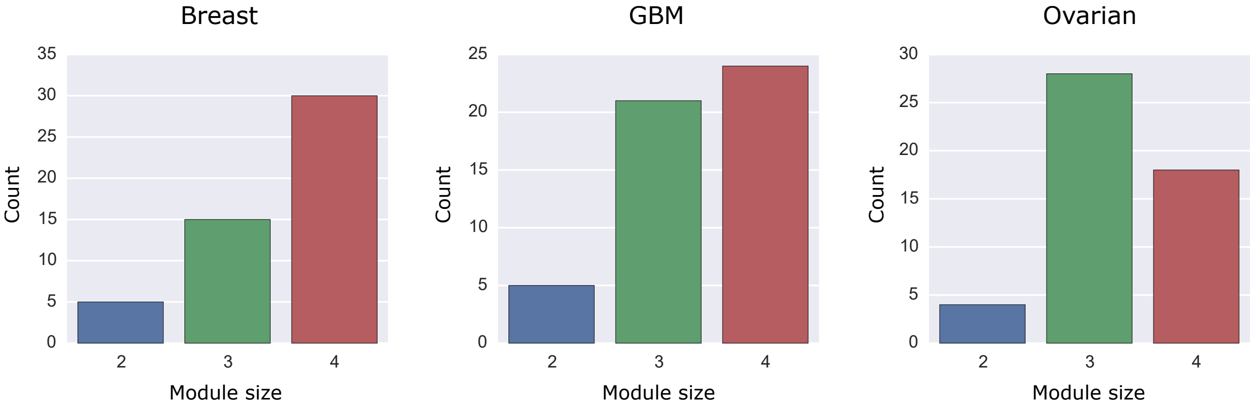

Figure S1: **Distribution of module sizes** as computed across the top 50 modules identified in breast cancer, GBM and ovarian cancer.

Table S1: Mean scores for the top 50 modules in breast cancer, by module size.

| Module size | ME | PPI | co-regulation | co-expression | ModulOmics |
|---|---|---|---|---|---|
| 2 | 0.92 | 0.71 | 0.40 | 0.55 | 0.64 |
| 3 | 0.87 | 0.82 | 0.64 | 0.30 | 0.66 |
| 4 | 0.77 | 0.95 | 0.49 | 0.23 | 0.61 |

Table S2: Mean scores for the top 50 modules in GBM, by module size.

| Module size | ME | PPI | co-regulation | co-expression | ModulOmics |
|---|---|---|---|---|---|
| 2 | 0.51 | 0.90 | 0.80 | 0.38 | 0.65 |
| 3 | 0.87 | 0.92 | 0.71 | 0.33 | 0.71 |
| 4 | 0.82 | 0.99 | 0.99 | 0.18 | 0.74 |

Table S3: Mean scores for top 50 modules in ovarian cancer, by module size.

| Module size | ME | PPI | co-regulation | co-expression | ModulOmics |
|---|---|---|---|---|---|
| 2 | 1.00 | 0.85 | 1.00 | 0.26 | 0.78 |
| 3 | 0.97 | 0.94 | 1.00 | 0.17 | 0.77 |
| 4 | 0.84 | 0.99 | 1.00 | 0.21 | 0.76 |

#### 3.2 Sensitivity analyses

We evaluated the sensitivity of the inferred modules to different parameter choices by running ModulOmics with changed parameter values and counting repetitions of results in the top 10 modules of size 4. We changed the parameters of the stochastic search as follows: 300 initial module seeds instead of the default 200, 15 clusters instead of the default 10, and 7 top results reported by each cluster instead of the default 5. We evaluated the following metrics: i) the repetition of gene connections, *i.e.* gene pairs co-residing in the same module, and ii) the repetition of the gene pool reported by the top modules, regardless of which module they belonged to. Across the three cancer types, the majority of gene pairs remained connected: 24, 19 and 16 gene pairs in breast cancer, GBM, and ovarian cancer, respectively, were common to the top modules inferred with each parameter configuration. Additionally, the majority of genes collectively included in the top modules were robust to parameter changes: 21, 8 and 8 genes respectively (Figure S2).

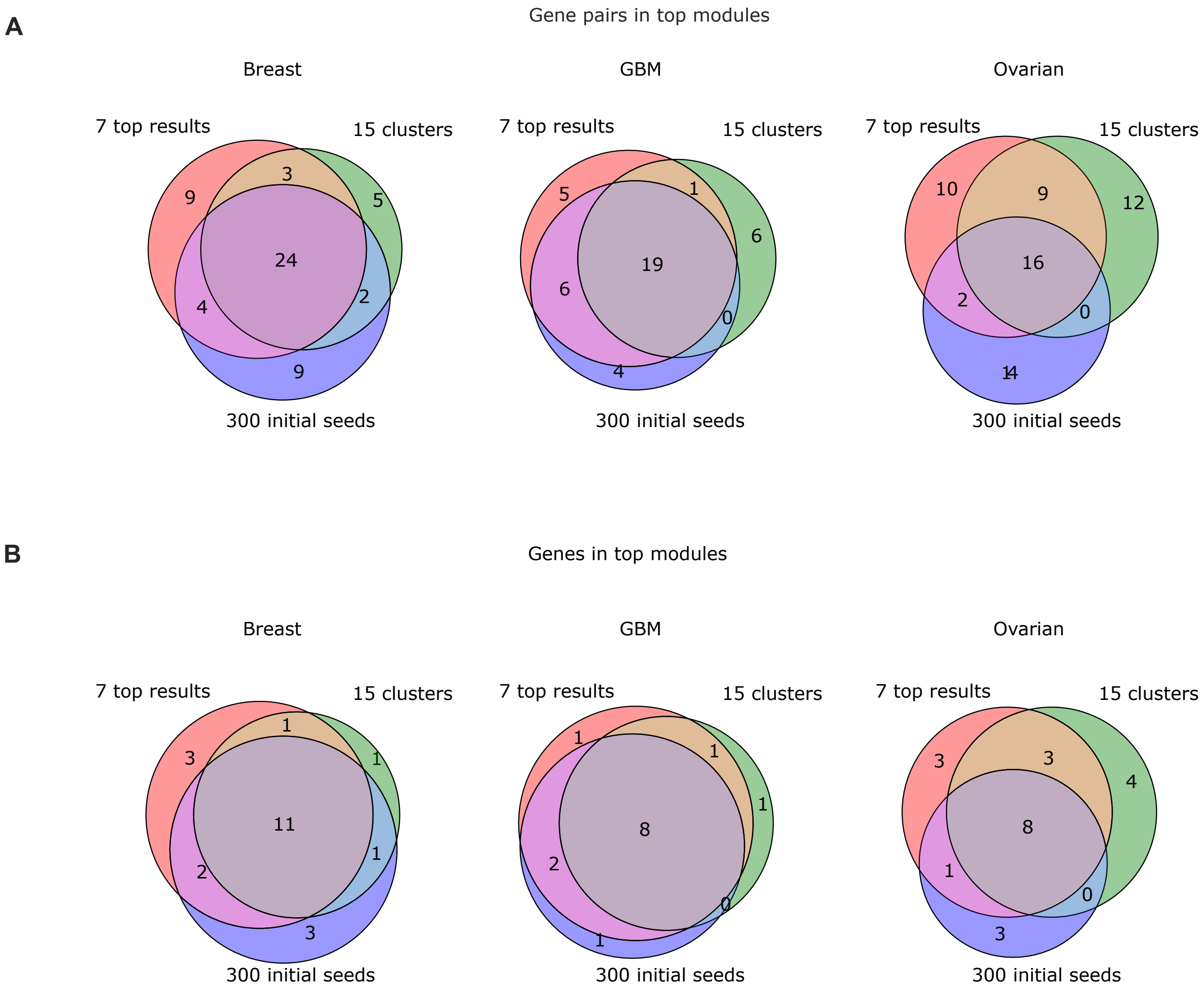

Figure S2: **Robustness of top 10 modules inferred by ModulOmics.** A) Intersection of gene pairs co-residing in the same module, for various parameter choices. B) Intersection of the gene pool collectively reported by the top 10 modules, for various parameter choices.

We further calculated how many suggested gene exchanges were accepted in the stochastic search (Table S4). As expected, for modules of size 2, no suggested exchanges were accepted, since the ILP solution recovers the pairwise global maximum. For modules of sizes 3 or 4, an average of 0.69 exchanges per module were accepted. The inferred modules were highly robust to increasing the number of suggested exchanges, such that using 30 exchanges yielded highly similar modules (data not shown).

Table S4: Gene exchanges made in the stochastic search, while refining the modules. The average # of accepted exchanges shown is per module.

| # exchanges | Cancer | Module size | Accepted exchanges | Rejected exchanges | Average # of accepted exchanges |
|---|---|---|---|---|---|
| 20 exchanges | Breast | 3 | 11 | 869 | 0.35 |
| 20 exchanges | Breast | 4 | 23 | 917 | 0.7 |
| 20 exchanges | GBM | 3 | 17 | 863 | 0.35 |
| 20 exchanges | GBM | 4 | 34 | 866 | 0.72 |
| 20 exchanges | Ovarian | 3 | 71 | 1209 | 1.39 |
| 20 exchanges | Ovarian | 4 | 93 | 1307 | 1.82 |

We next evaluated the contribution of the stochastic search to the functional connectivity of the resulted modules by exploring the properties of their enrichment with known pathways (see details about pathway enrichment calculation in section 3.4). To this end, we compared the mean number of module members in an enriched pathway, as well as the mean enrichment score, before and after the stochastic search (Figure S3). We concentrated on the top 30 modules of largest size (size 4, with the highest rate of accepted exchanges during the stochastic search, as demonstrated by Table S4). Reassuringly, both metrics showed that the stochastic search improves the identification of functionally connected groups.

Table S5 shows the fixed parameters used by ModulOmics in computing the individual scores, as well as in the stochastic search.

Table S5: Fixed parameters used by ModulOmics.

| Context | Parameter | Default value | Explanation |
|---|---|---|---|
| Co-Expression test | Expression quantile | k = 2<br>q = 4 | A gene is defined as expressed if its expression averaged across all samples is above the $k^{\text{th}}$ $q$-quantile. Genes that are unexpressed are also not co-expressed. |
| Co-regulation test | active TF | $\lvert zScore\rvert > 1$<br>in 25% of the samples | Active TF in the cohort |
| Module search | Initial seeds | 300 | Number of initial modules inferred by the ILP based on pairwise scores. These seeds are then further refined by the stochastic search to maximize the global ModulOmics score. |
| Module search | Number of clusters | 10 | Number of clusters in which the seed modules are divided, so as to start the search each time from a different cluster. |
| Module search | Top results | 5 | Top modules reported by each cluster. The top modules are aggregated to one list which is re-ranked and retrieved to the user. |

In this section, we showed that the modules identified by ModulOmics are highly robust to changes in the module search parameters. To choose default parameter values for the co-regulation and co-expression tests

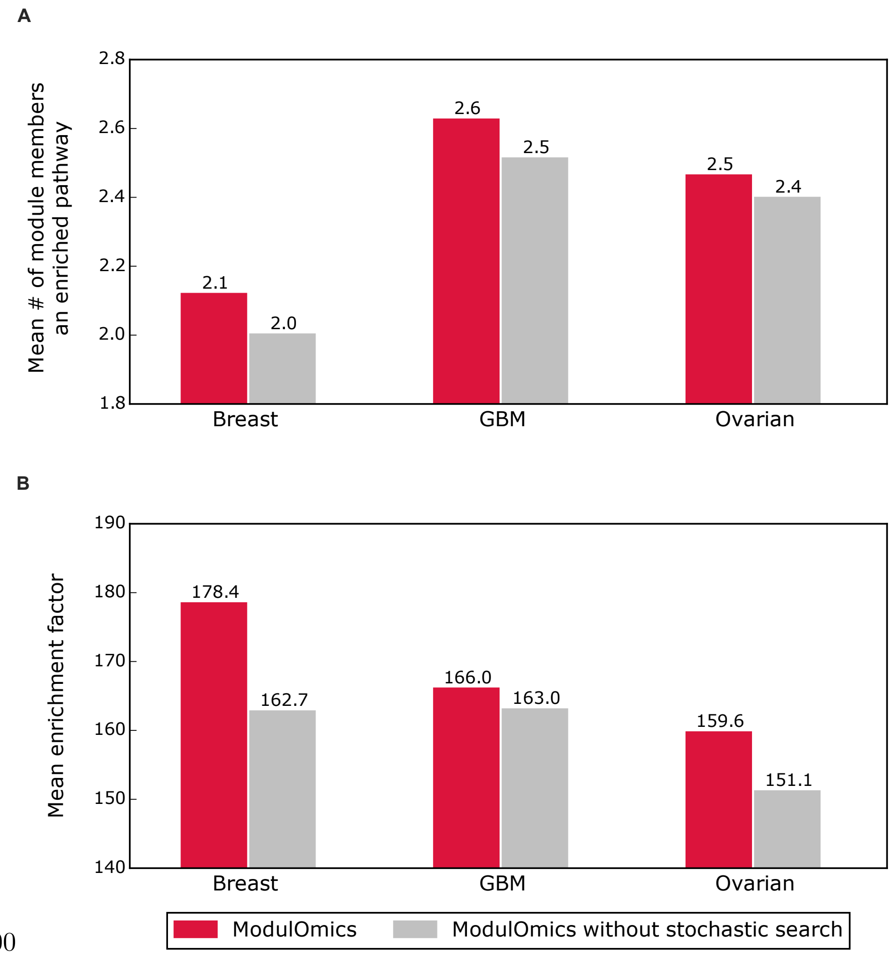

000

Figure S3: **Pathway enrichment analysis for ModulOmics vs. ModulOmics without using the stochastic search for the top** 30 **modules of size** 4. A) Mean number of genes in a module to participate in a significantly enriched pathway. B) Mean enrichment factor score per module.

we scanned a range of possible values, and chose values which included a substantial amount of information for all the studied cohorts. For example, the default values chosen for the co-regulation test yielded for GBM, breast and ovarian cancer 204, 195 and 232 active transcription factors respectively. Increasing the zScore threshold to 2 (as traditionally used by TCGA) yielded only 1, 2 and 9 respectively, substantially reducing the information retrieved. Similarly, the default parameter value for the co-expression test, namely calling a gene as expressed in the cohort if its expression averaged across all samples was above the $2^{nd}$ $4^{th}$-quantile, yielded for GBM, breast and ovarian cancer 14,823, 9,264 and 4,064 comparable pairs respectively (out of 263,682, 79,401 and 33,153 potential gene pairs). Making the threshold more stringent by setting it to the $3^{rd}$ $4^{th}$-quantile yielded only 3,690,2,328 and 1,008 comparable pairs.

Having explored the values for the three different cohorts, we recommend the default values described in Table S5, yet encourage the user to test a range of values when running ModulOmics on a new cohort.

#### 3.3 Driver modules are enriched with cancer drivers

##### 3.3.1 Positive and negative control lists

To evaluate the enrichment of modules with known driver genes, we used a large number of control lists, derived from independent sources. As positive controls, we used the following lists, introduced in [10]: i) The Cancer Gene Census (CGC) version 73 (*PosSomatic* and *PosTrans*), a set of 569 genes manually curated by The Sanger Institute, which have alterations in somatic and germline SNVs, CNVs and translocations; ii) UniprotKB [22] (*PosUniprotKB*), a manually curated database of 412 functional proteins, classified as proto-oncogene, oncogene and tumour suppressor gene; iii) a query of DISEASES [15] (*PosTextMine*), a database of disease-gene associations extracted mainly from text-mining, which consists of 711 genes associated with cancer; and iv) The Atlas of Genetics and Cytogenetics in Oncology and Hematology (*PosAGO*) [11], a list of $1,430$ cancer genes manually curated by a collaborative effort spanning multiple centers. *PosUnionAll* is the union of all these positive control lists. In addition, we used the Network of Cancer Genes (NCG5) [1], a manually curated list consisting of $1,571$ protein-coding cancer driver genes compiled by The Sanger Institute. The gene members of the two shortest CGC lists, germline SNVs (38 genes) and CNVs (15 genes), were not identified in any high scoring module by any of the tested methods, hence are not shown here.

To evaluate the enrichment of modules with known non-driver genes, we used the following lists introduced in [10], as negative controls: i) a list derived from AGO [11] consisting of $9,457$ genes that have no evidence of association with cancer (*NegAgoFull*); ii) a conservative version of the negative AGO list (*NegAGOClean*), created by filtering genes that are part of any cancer-related pathway from the MSigDB database [19], resulting in $3,272$ genes, and iii) a list of known non-driver genes introduced in [7] (*NegDavoli*).

##### 3.3.2 Driver enrichment in top modules

According to the enrichment of the highest scoring driver modules with known driver genes (positive controls) and known non-driver genes (negative controls), ModulOmics generally outperformed the three competitive methods evaluated: TiMEx [6], HotNet2 [13], and MEMCover [12], as well as the four single-omics and the four three-omics approaches, for groups of sizes 2, 3 and 4 (Figures S4 and S5). MEMCover identified only 3 modules of size 3 in breast cancer, and 1 module in GBM and ovarian cancer, as well as only 1 module of size 4 in any of the three cancer types; TiMEx identified only 3 modules of size 3 in ovarian cancer, 1 module of size 4 in breast cancer, and no modules in GBM or ovarian cancer; HotNet2 identified only 10 modules of size 2 in breast cancer, 8 modules in GBM, and 2 modules in ovarian cancer, only 5 modules of size 3 in breast cancer, 2 modules in GBM, and no modules in ovarian, as well as no modules of size 4 in any of the three cancer types. These performances per module size are therefore not shown in Figure S4. Table S6 displays the scores plotted in Figures 2, S4 and S5B.

##### 3.3.3 Driver genes enrichment in top unique genes

One of the features of ModulOmics is that each gene can participate in multiple modules (Figure S6). ModulOmics outperformed the other methods also when restricting the comparison to the same number of unique genes part of the top modules, regardless of their membership (Figure S9).

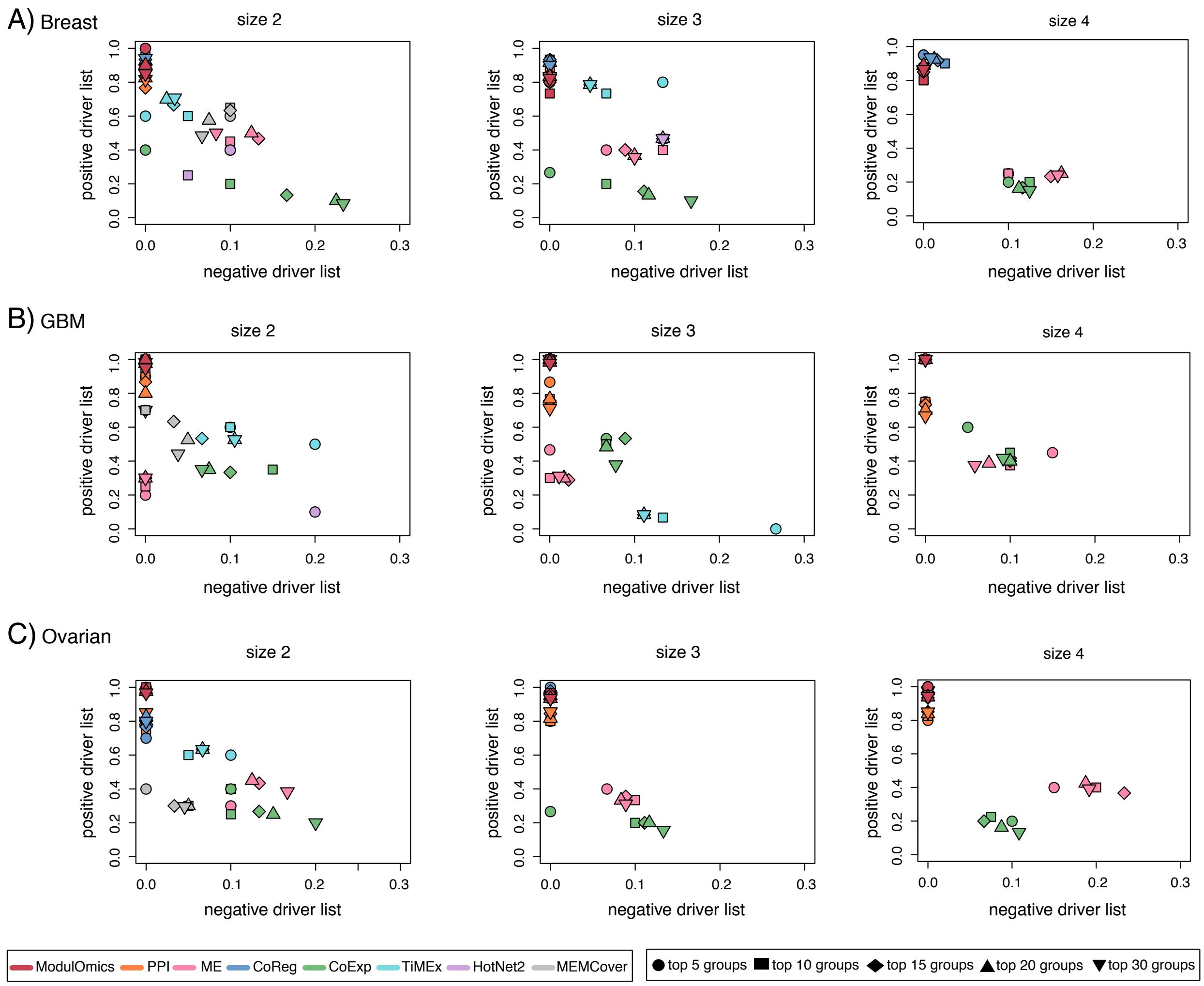

Figure S4: **The driver modules inferred by ModulOmics are enriched with cancer driver genes (extending Figure 2B in the main text).** Detailed driver and non-driver enrichment scores for the positive driver list *PosUnionAll* and the negative driver list *NegAGOClean* for the eight methods assessed for the top scoring 5, 10, 15, 20 and 30 modules and for the cancer types **A)** breast, **B)** GBM and **C)** ovarian.

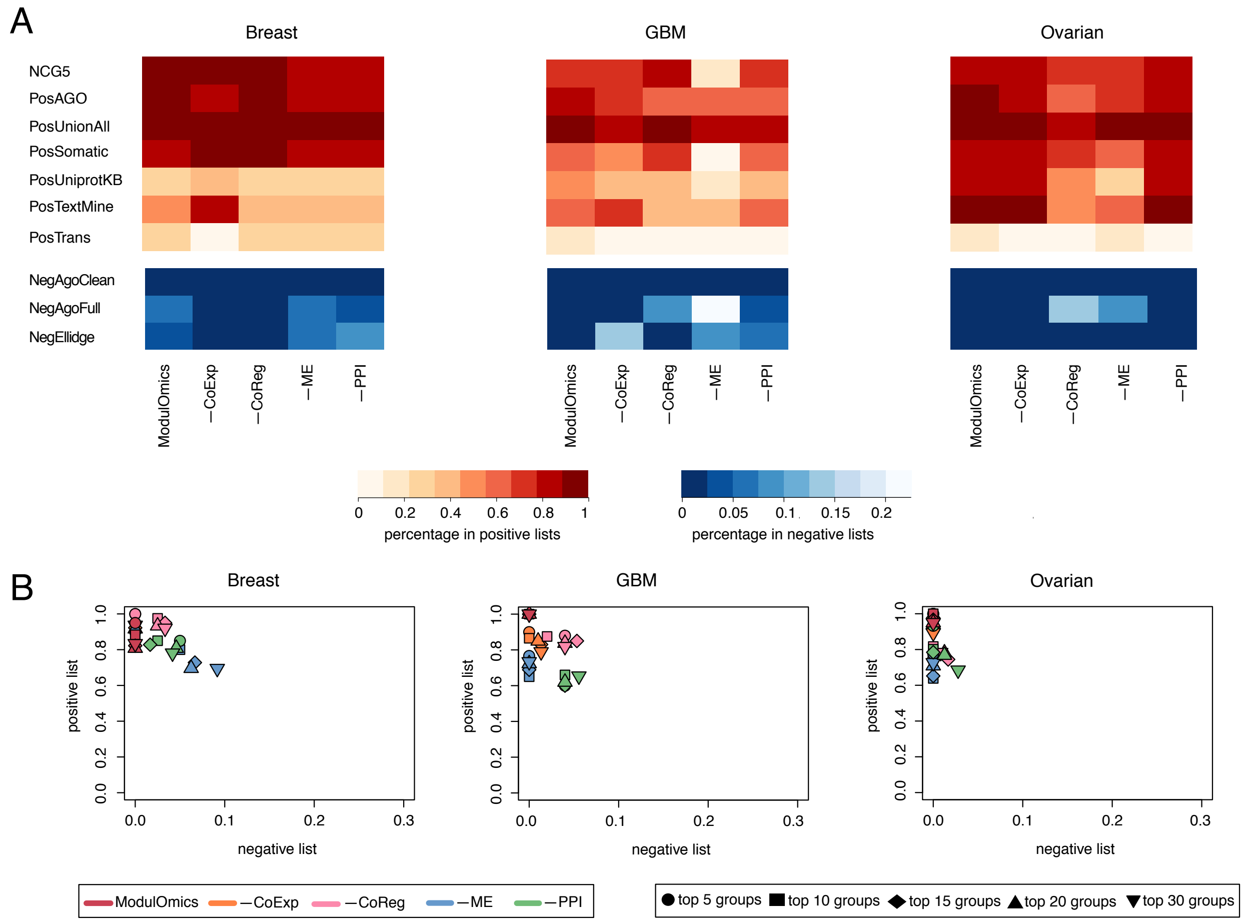

Figure S5: **The driver modules inferred by ModulOmics are enriched with cancer driver genes (extending Figure 2 in the main text). A)** The average driver enrichment (red heatmaps) and non-driver enrichment (blue heatmaps) across the top 10 scoring modules inferred by ModulOmics and by using only subsets of three omics data sources. **B)** Detailed driver and non-driver enrichment scores for the positive driver list *PosUnionAll* and the negative driver list *NegAGOClean* for ModulOmics and the subsets of three omics for the top scoring 5, 10, 15, 20 and 30 modules. The − sign next to an omic data source represents the situation in which that omic data source was excluded from the computation of the ModulOmics score.

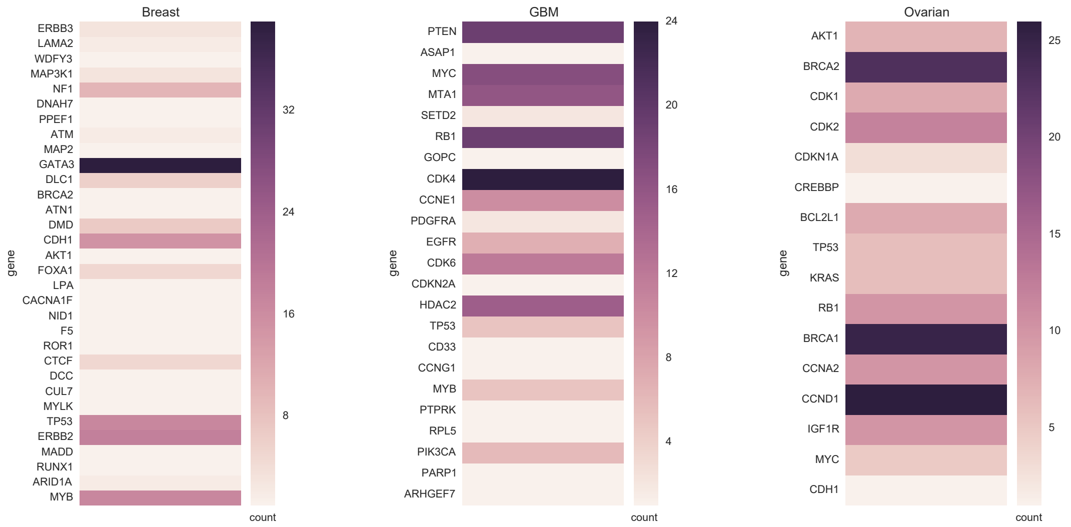

Figure S6: **Occurrence of genes in the top 50 modules identified by ModulOmics across the three cancer types.**

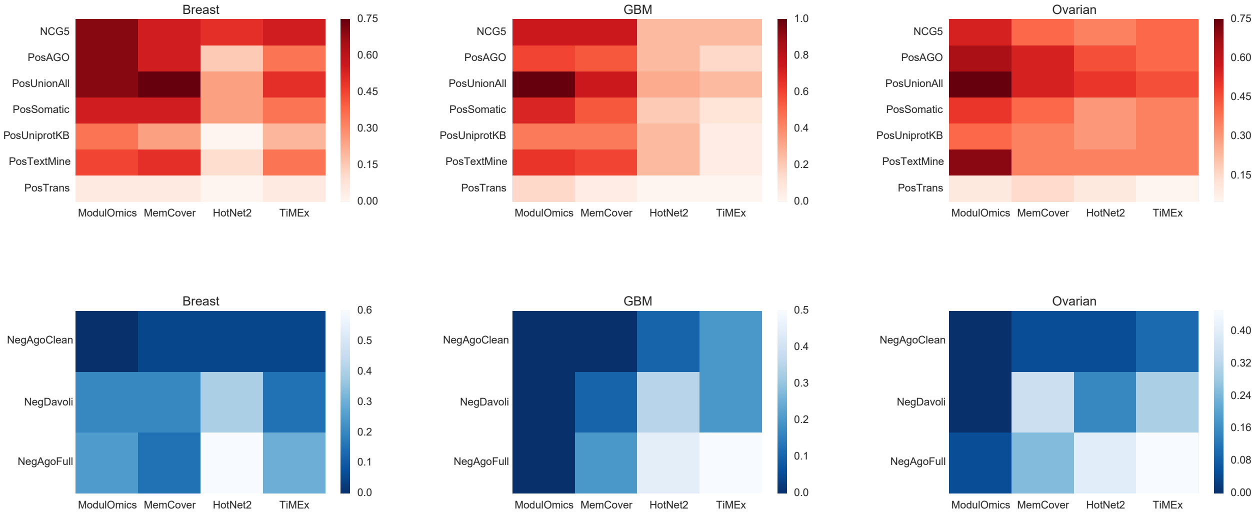

Figure S7: **The 20 unique genes in the top driver modules inferred by ModulOmics are more enriched with cancer driver genes.**

Table S6: Scores corresponding to the performance of the methods displayed in Figures 2B, S4 and S5B. Each pair $(a, b)$ represents the average driver enrichment scores for the positive control list $PosUnionAll$ and the negative control list $NegAGOClean$.

| Figure | Size | Method | Top | Breast cancer | GBM | Ovarian cancer |
|---|---|---|---|---|---|---|
| 2B | ALL | ModulOmics | 5 | $(0.95, 0)$ | $(1, 0)$ | $(1, 0)$ |
| 2B | ALL | ModulOmics | 10 | $(0.83, 0)$ | $(1, 0)$ | $(1, 0)$ |
| 2B | ALL | ModulOmics | 15 | $(0.82, 0)$ | $(1, 0)$ | $(0.96, 0)$ |
| 2B | ALL | ModulOmics | 20 | $(0.80, 0)$ | $(1, 0)$ | $(0.95, 0)$ |
| 2B | ALL | ModulOmics | 30 | $(0.83, 0)$ | $(1, 0)$ | $(0.95, 0)$ |
| S4 | 2 | ModulOmics | 5 | $(1, 0)$ | $(1, 0)$ | $(1, 0)$ |
| S4 | 2 | ModulOmics | 10 | $(0.85, 0)$ | $(1, 0)$ | $(1, 0)$ |
| S4 | 2 | ModulOmics | 15 | $(0.9, 0)$ | $(1, 0)$ | $(0.96, 0)$ |
| S4 | 2 | ModulOmics | 20 | $(0.9, 0)$ | $(1, 0)$ | $(0.97, 0)$ |
| S4 | 2 | ModulOmics | 30 | $(0.85, 0)$ | $(0.95, 0)$ | $(0.96, 0)$ |
| S4 | 3 | ModulOmics | 5 | $(0.8, 0)$ | $(1, 0)$ | $(0.93, 0)$ |
| S4 | 3 | ModulOmics | 10 | $(0.73, 0)$ | $(1, 0)$ | $(0.96, 0)$ |
| S4 | 3 | ModulOmics | 15 | $(0.8, 0)$ | $(1, 0)$ | $(0.95, 0)$ |
| S4 | 3 | ModulOmics | 20 | $(0.83, 0)$ | $(0.98, 0)$ | $(0.93, 0)$ |
| S4 | 3 | ModulOmics | 30 | $(0.82, 0)$ | $(0.97, 0)$ | $(0.93, 0)$ |
| S4 | 4 | ModulOmics | 5 | $(0.85, 0)$ | $(1, 0)$ | $(1, 0)$ |
| S4 | 4 | ModulOmics | 10 | $(0.8, 0)$ | $(1, 0)$ | $(0.95, 0)$ |
| S4 | 4 | ModulOmics | 15 | $(0.85, 0)$ | $(1, 0)$ | $(0.93, 0)$ |
| S4 | 4 | ModulOmics | 20 | $(0.88, 0)$ | $(1, 0)$ | $(0.93, 0)$ |
| S4 | 4 | ModulOmics | 30 | $(0.86, 0)$ | $(1, 0)$ | $(0.94, 0)$ |
| 2B | ALL | PPI | 5 | $(0.8, 0)$ | $(0.93, 0)$ | $(0.78, 0)$ |
| 2B | ALL | PPI | 10 | $(0.85, 0)$ | $(0.84, 0)$ | $(0.79, 0)$ |
| 2B | ALL | PPI | 15 | $(0.87, 0)$ | $(0.79, 0)$ | $(0.79, 0)$ |
| 2B | ALL | PPI | 20 | $(0.87, 0)$ | $(0.8, 0)$ | $(0.79, 0)$ |
| 2B | ALL | PPI | 30 | $(0.83, 0)$ | $(0.81, 0)$ | $(0.79, 0)$ |
| S4 | 2 | PPI | 5 | $(0.9, 0)$ | $(0.9, 0)$ | $(0.8, 0)$ |
| S4 | 2 | PPI | 10 | $(0.85, 0)$ | $(0.9, 0)$ | $(0.75, 0)$ |
| S4 | 2 | PPI | 15 | $(0.76, 0)$ | $(0.86, 0)$ | $(0.8, 0)$ |
| S4 | 2 | PPI | 20 | $(0.82, 0)$ | $(0.8, 0)$ | $(0.8, 0)$ |
| S4 | 2 | PPI | 30 | $(0.81, 0)$ | $(0.7, 0)$ | $(0.85, 0)$ |
| S4 | 3 | PPI | 5 | $(0.8, 0)$ | $(0.86, 0)$ | $(0.8, 0)$ |
| S4 | 3 | PPI | 10 | $(0.86, 0)$ | $(0.76, 0)$ | $(0.8, 0)$ |
| S4 | 3 | PPI | 15 | $(0.84, 0)$ | $(0.75, 0)$ | $(0.84, 0)$ |
| S4 | 3 | PPI | 20 | $(0.83, 0)$ | $(0.76, 0)$ | $(0.81, 0)$ |
| S4 | 3 | PPI | 30 | $(0.83, 0)$ | $(0.71, 0)$ | $(0.85, 0)$ |
| S4 | 4 | PPI | 5 | $(0.95, 0)$ | $(0.75, 0)$ | $(0.8, 0)$ |
| S4 | 4 | PPI | 10 | $(0.87, 0)$ | $(0.75, 0)$ | $(0.82, 0)$ |
| S4 | 4 | PPI | 15 | $(0.88, 0)$ | $(0.73, 0)$ | $(0.85, 0)$ |
| S4 | 4 | PPI | 20 | $(0.88, 0)$ | $(0.7, 0)$ | $(0.83, 0)$ |
| S4 | 4 | PPI | 30 | $(0.85, 0)$ | $(0.66, 0)$ | $(0.85, 0)$ |
| 2B | ALL | ME | 5 | $(0.4, 0.06)$ | $(0.2, 0)$ | $(0.4, 0.06)$ |
| 2B | ALL | ME | 10 | $(0.4, 0.13)$ | $(0.25, 0)$ | $(0.33, 0.1)$ |
| 2B | ALL | ME | 15 | $(0.4, 0.08)$ | $(0.3, 0)$ | $(0.35, 0.08)$ |
| 2B | ALL | ME | 20 | $(0.36, 0.1)$ | $(0.3, 0)$ | $(0.33, 0.08)$ |
| 2B | ALL | ME | 30 | $(0.35, 0.1)$ | $(0.3, 0)$ | $(0.31, 0.08)$ |
| S4 | 2 | ME | 5 | $(0.4, 0.1)$ | $(0.2, 0)$ | $(0.3, 0.1)$ |
| S4 | 2 | ME | 10 | $(0.45, 0.1)$ | $(0.25, 0)$ | $(0.4, 0.1)$ |
| S4 | 2 | ME | 15 | $(0.46, 0.13)$ | $(0.3, 0)$ | $(0.43, 0.13)$ |
| S4 | 2 | ME | 20 | $(0.5, 0.12)$ | $(0.3, 0)$ | $(0.45, 0.12$ |

| | | | | | | |
|---|---|---|---|---|---|---|
| S4 | 2 | ME | 30 | $(0.5, 0.08)$ | $(0.3, 0)$ | $(0.38, 0.16)$ |
| S4 | 3 | ME | 5 | $(0.4, 0.06)$ | $(0.46, 0)$ | $(0.4, 0.06)$ |
| S4 | 3 | ME | 10 | $(0.4, 0.13)$ | $(0.3, 0)$ | $(0.33, 0.1)$ |
| S4 | 3 | ME | 15 | $(0.4, 0.08)$ | $(0.28, 0.02)$ | $(0.35, 0.08)$ |
| S4 | 3 | ME | 20 | $(0.36, 0.1)$ | $(0.3, 0.01)$ | $(0.33, 0.08$ |
| S4 | 3 | ME | 30 | $(0.35, 0.1)$ | $(0.31, 0.01)$ | $(0.31, 0.08)$ |
| S4 | 4 | ME | 5 | $(0.25, 0.1)$ | $(0.45, 0.15)$ | $(0.4, 0.15)$ |
| S4 | 4 | ME | 10 | $(0.25, 0.1)$ | $(0.37, 0.1)$ | $(0.4, 0.2)$ |
| S4 | 4 | ME | 15 | $(0.23, 0.15)$ | $(0.41, 0.1)$ | $(0.36, 0.23)$ |
| S4 | 4 | ME | 20 | $(0.25, 0.16)$ | $(0.38, 0.07)$ | $(0.42, 0.18$ |
| S4 | 4 | ME | 30 | $(0.24, 0.15)$ | $(0.37, 0.05)$ | $(0.39, 0.19)$ |
| 2B | ALL | CoReg | 5 | $(0.93, 0)$ | $(1, 0)$ | $(0.7, 0)$ |
| 2B | ALL | CoReg | 10 | $(0.93, 0)$ | $(0.95, 0)$ | $(0.8, 0)$ |
| 2B | ALL | CoReg | 15 | $(0.93, 0)$ | $(0.96, 0)$ | $(0.76, 0)$ |
| 2B | ALL | CoReg | 20 | $(0.95, 0)$ | $(0.97, 0)$ | $(0.82, 0)$ |
| 2B | ALL | CoReg | 30 | $(0.95, 0)$ | $(0.98, 0)$ | $(0.8, 0)$ |
| S4 | 2 | CoReg | 5 | $(1, 0)$ | $(1, 0)$ | $(0.7, 0)$ |
| S4 | 2 | CoReg | 10 | $(0.95, 0)$ | $(0.95, 0)$ | $(0.8, 0)$ |
| S4 | 2 | CoReg | 15 | $(0.93, 0)$ | $(0.96, 0)$ | $(0.76, 0)$ |
| S4 | 2 | CoReg | 20 | $(0.92, 0)$ | $(0.97, 0)$ | $(0.82, 0)$ |
| S4 | 2 | CoReg | 30 | $(0.94, 0)$ | $(0.98, 0)$ | $(0.8, 0)$ |
| S4 | 3 | CoReg | 5 | $(0.93, 0)$ | $(1, 0)$ | $(1, 0)$ |
| S4 | 3 | CoReg | 10 | $(0.93, 0)$ | $(1, 0)$ | $(0.96, 0)$ |
| S4 | 3 | CoReg | 15 | $(0.93, 0)$ | $(1, 0)$ | $(0.97, 0)$ |
| S4 | 3 | CoReg | 20 | $(0.91, 0)$ | $(1, 0)$ | $(0.95, 0)$ |
| S4 | 3 | CoReg | 30 | $(0.9, 0)$ | $(1, 0)$ | $(0.94, 0)$ |
| S4 | 4 | CoReg | 5 | $(0.95, 0)$ | $(1, 0)$ | $(1, 0)$ |
| S4 | 4 | CoReg | 10 | $(0.9, 0.025)$ | $(1, 0)$ | $(0.97, 0)$ |
| S4 | 4 | CoReg | 15 | $(0.91, 0.01)$ | $(1, 0)$ | $(0.96, 0)$ |
| S4 | 4 | CoReg | 20 | $(0.92, 0.01)$ | $(1, 0)$ | $(0.97, 0)$ |
| S4 | 4 | CoReg | 30 | $(0.93, 0.08)$ | $(1, 0)$ | $(0.97, 0)$ |
| 2B | ALL | CoExp | 5 | $(0.38, 0)$ | $(0.5, 0.16)$ | $(0.38, 0)$ |
| 2B | ALL | CoExp | 10 | $(0.32, 0.05)$ | $(0.51, 0.1)$ | $(0.32, 0.05)$ |
| 2B | ALL | CoExp | 15 | $(0.23, 0.06)$ | $(0.55, 0.07)$ | $(0.28, 0.05)$ |
| 2B | ALL | CoExp | 20 | $(0.21, 0.07)$ | $(0.47, 0.1)$ | $(0.24, 0.07)$ |
| 2B | ALL | CoExp | 30 | $(0.19, 0.12)$ | $(0.43, 0.1)$ | $(0.23, 0.1)$ |
| S4 | 2 | CoExp | 5 | $(0.4, 0)$ | $(0.6, 0.1)$ | $(0.4, 0.1)$ |
| S4 | 2 | CoExp | 10 | $(0.2, 0.1)$ | $(0.35, 0.15)$ | $(0.25, 0.1)$ |
| S4 | 2 | CoExp | 15 | $(0.13, 0.16)$ | $(0.33, 0.1)$ | $(0.26, 0.13)$ |
| S4 | 2 | CoExp | 20 | $(0.1, 0.22)$ | $(0.35, 0.075)$ | $(0.25, 0.15)$ |
| S4 | 2 | CoExp | 30 | $(0.08, 0.23)$ | $(0.35, 0.06)$ | $(0.2, 0.2)$ |
| S4 | 3 | CoExp | 5 | $(0.26, 0)$ | $(0.53, 0.06)$ | $(0.26, 0)$ |
| S4 | 3 | CoExp | 10 | $(0.2, 0.06)$ | $(0.5, 0.06)$ | $(0.15, 0.13)$ |
| S4 | 3 | CoExp | 15 | $(0.15, 0.11)$ | $(0.53, 0.08)$ | $(0.2, 0.11)$ |
| S4 | 3 | CoExp | 20 | $(0.13, 0.11)$ | $(0.48, 0.06)$ | $(0.2, 0.11)$ |
| S4 | 3 | CoExp | 30 | $(0.1, 0.16)$ | $(0.37, 0.07)$ | $(0.2, 0.1)$ |
| S4 | 4 | CoExp | 5 | $(0.2, 0.1)$ | $(0.6, 0.05)$ | $(0.2, 0.1)$ |
| S4 | 4 | CoExp | 10 | $(0.2, 0.12)$ | $(0.45, 0.1)$ | $(0.22, 0.07)$ |
| S4 | 4 | CoExp | 15 | $(0.16, 0.11)$ | $(0.4, 0.1)$ | $(0.2, 0.06)$ |
| S4 | 4 | CoExp | 20 | $(0.16, 0.11)$ | $(0.4, 0.1)$ | $(0.16, 0.08)$ |
| S4 | 4 | CoExp | 30 | $(0.15, 0.12)$ | $(0.41, 0.09)$ | $(0.13, 0.1)$ |
| 2B | ALL | MEMCover | 5 | $(0.6, 0.1)$ | $(0.7, 0)$ | $(0.55, 0)$ |
| 2B | ALL | MEMCover | 10 | $(0.75, 0.05)$ | $(0.7, 0)$ | $(0.42, 0.05)$ |
| 2B | ALL | MEMCover | 15 | $(0.65, 0.06)$ | $(0.7, 0.03)$ | $(0.35, 0.03)$ |

| | | | | | | |
|---|---|---|---|---|---|---|
| 2B | ALL | MEMCover | 20 | $(0.69, 0.07)$ | $(0.57, 0.05)$ | $(0.36, 0.05)$ |
| 2B | ALL | MEMCover | 30 | $(0.54, 0.06)$ | $(0.46, 0.03)$ | $(0.34, 0.04)$ |
| S4 | 2 | MEMCover | 5 | $(0.6, 0.1)$ | $(0.7, 0)$ | $(0.4, 0)$ |
| S4 | 2 | MEMCover | 10 | $(0.65, 0.1)$ | $(0.7, 0)$ | $(0.3, 0.05)$ |
| S4 | 2 | MEMCover | 15 | $(0.63, 0.06)$ | $(0.63, 0.03)$ | $(0.3, 0.03)$ |
| S4 | 2 | MEMCover | 20 | $(0.57, 0.07)$ | $(0.52, 0.05)$ | $(0.3, 0.05)$ |
| S4 | 2 | MEMCover | 30 | $(0.48, 0.06)$ | $(0.44, 0.03)$ | $(0.29, 0.04)$ |
| 2B | ALL | TiMEx | 5 | $(0.6, 0)$ | $(0.5, 0.2)$ | $(0.6, 0.1)$ |
| 2B | ALL | TiMEx | 10 | $(0.6, 0.05)$ | $(0.6, 0.1)$ | $(0.6, 0.05)$ |
| 2B | ALL | TiMEx | 15 | $(0.66, 0.03)$ | $(0.4, 0.15)$ | $(0.67, 0.05)$ |
| 2B | ALL | TiMEx | 20 | $(0.7, 0.025)$ | $(0.31, 0.11)$ | $(0.65, 0.09)$ |
| 2B | ALL | TiMEx | 30 | $(0.73, 0.03)$ | $(0.35, 0.09)$ | $(0.65, 0.09)$ |
| S4 | 2 | TiMEx | 5 | $(0.6, 0)$ | $(0.5, 0.2)$ | $(0.6, 0.1)$ |
| S4 | 2 | TiMEx | 10 | $(0.6, 0.05)$ | $(0.6, 0.1)$ | $(0.6, 0.05)$ |
| S4 | 2 | TiMEx | 15 | $(0.66, 0.03)$ | $(0.53, 0.06)$ | $(0.63, 0.06)$ |
| S4 | 2 | TiMEx | 20 | $(0.7, 0.025)$ | $(0.52, 0.1)$ | $(0.63, 0.06)$ |
| S4 | 2 | TiMEx | 30 | $(0.7, 0.03)$ | $(0.52, 0.1)$ | $(0.63, 0.06)$ |
| S4 | 3 | TiMEx | 5 | $(0.8, 0.13)$ | $(0, 0.26)$ | |
| S4 | 3 | TiMEx | 10 | $(0.73, 0.06)$ | $(0.06, 0.13)$ | |
| S4 | 3 | TiMEx | 15 | $(0.78, 0.04)$ | $(0.08, 0.11)$ | |
| S4 | 3 | TiMEx | 20 | $(0.78, 0.04)$ | $(0.08, 0.11)$ | |
| S4 | 3 | TiMEx | 30 | $(0.78, 0.04)$ | $(0.08, 0.11)$ | |
| 2B | ALL | HotNet2 | 5 | $(0.42, 0.09)$ | $(0.71, 0)$ | $(0.53, 0.1)$ |
| 2B | ALL | HotNet2 | 10 | $(0.42, 0.11)$ | $(0.51, 0.05)$ | $(0.44, 0.09)$ |
| 2B | ALL | HotNet2 | 15 | $(0.47, 0.1)$ | $(0.37, 0.06)$ | $(0.44, 0.09)$ |
| 2B | ALL | HotNet2 | 20 | $(0.4, 0.08)$ | $(0.41, 0.06)$ | $(0.44, 0.09)$ |
| 2B | ALL | HotNet2 | 30 | $(0.36, 0.07)$ | $(0.41, 0.06)$ | $(0.44, 0.09)$ |
| S4 | 2 | HotNet2 | 5 | $(0.4, 0.1)$ | $(0.1, 0.2)$ | |
| S4 | 2 | HotNet2 | 10 | $(0.25, 0.05)$ | | |
| S4 | 3 | HotNet2 | 5 | $(0.46, 0.13)$ | | |
| S5B | ALL | -CoExp | 5 | $(1, 0)$ | $(0.9, 0)$ | $(1, 0)$ |
| S5B | ALL | -CoExp | 10 | $(0.93, 0)$ | $(0.86, 0)$ | $(0.96, 0)$ |
| S5B | ALL | -CoExp | 15 | $(0.92, 0)$ | $(0.83, 0.01)$ | $(0.97, 0)$ |
| S5B | ALL | -CoExp | 20 | $(0.91, 0)$ | $(0.85, 0.01)$ | $(0.93, 0)$ |
| S5B | ALL | -CoExp | 30 | $(0.93, 0)$ | $(0.78, 0.01)$ | $(0.89, 0)$ |
| S5B | ALL | -CoReg | 5 | $(1, 0)$ | $(0.88, 0.04)$ | $(0.75, 0)$ |
| S5B | ALL | -CoReg | 10 | $(0.97, 0.02)$ | $(0.87, 0.02)$ | $(0.81, 0)$ |
| S5B | ALL | -CoReg | 15 | $(0.95, 0.03)$ | $(0.85, 0.05)$ | $(0.74, 0.01)$ |
| S5B | ALL | -CoReg | 20 | $(0.93, 0.02)$ | $(0.83, 0.04)$ | $(0.78, 0.01)$ |
| S5B | ALL | -CoReg | 30 | $(0.91, 0.03)$ | $(0.81, 0.04)$ | $(0.78, 0)$ |
| S5B | ALL | -ME | 5 | $(0.9, 0)$ | $(0.76, 0.04)$ | $(0.77, 0)$ |
| S5B | ALL | -ME | 10 | $(0.8, 0.05)$ | $(0.65, 0.02)$ | $(0.63, 0)$ |
| S5B | ALL | -ME | 15 | $(0.72, 0.06)$ | $(0.68, 0.05)$ | $(0.65, 0.01)$ |
| S5B | ALL | -ME | 20 | $(0.69, 0.06)$ | $(0.72, 0.04)$ | $(0.7, 0.01)$ |
| S5B | ALL | -ME | 30 | $(0.69, 0.09)$ | $(0.73, 0.04)$ | $(0.72, 0)$ |
| S5B | ALL | -PPI | 5 | $(0.85, 0.05)$ | $(0.6, 0.04)$ | $(0.93, 0)$ |
| S5B | ALL | -PPI | 10 | $(0.85, 0.02)$ | $(0.66, 0.04)$ | $(0.8, 0)$ |
| S5B | ALL | -PPI | 15 | $(0.82, 0.01)$ | $(0.6, 0.04)$ | $(0.78, 0.01)$ |
| S5B | ALL | -PPI | 20 | $(0.8, 0.04)$ | $(0.62, 0.04)$ | $(0.76, 0.01)$ |
| S5B | ALL | -PPI | 30 | $(0.78, 0.04)$ | $(0.65, 0.05)$ | $(0.68, 0)$ |

##### 3.4 Driver modules are functionally coherent

To evaluate the enrichment of modules with known pathways, we used two statistical tests of module-pathway intersection, as proposed by the Expander software [21]: i) a hyper-geometric enrichment test to calculate the occurrence probability of the intersection of a module with a pathway at random when drawing from all gene coding proteins, and ii) an enrichment factor designed to ease the bias towards larger modules. The enrichment factor is defined as the ratio between the sizes of the intersection of each module and each pathway and the intersection of that pathway and the set of all background genes (all protein coding genes), normalized by the sizes of the module and background respectively: $\frac{|\text{module} \cap \text{pathway}|}{|\text{pathway} \cap \text{background genes}|} \times \frac{|\text{background genes}|}{|\text{module}|}$

###### 3.4.1 Pathways per module size

Analyzing the pathway enrichment per module size, Figure S8 shows that no particular size dominates the enriched pathways, with the exception of modules of size 2 in ovarian cancer, which are less enriched with known pathways.

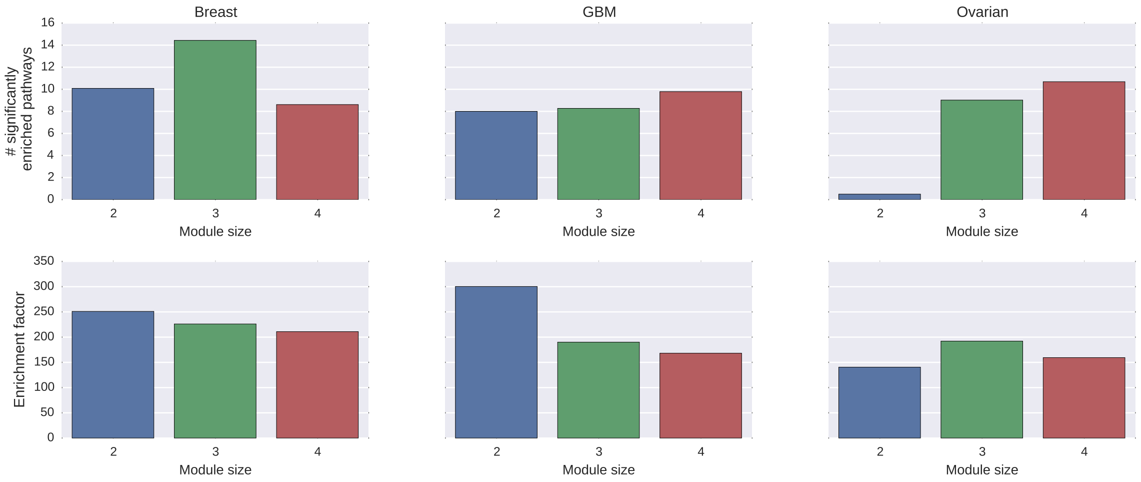

Figure S8: **Pathway enrichment per module size** based on the 30 top modules identified by ModulOmics.

###### 3.4.2 Pathway enrichment in subsets of three omics

We further evaluated the contribution of each single omic to pathway enrichment, by running ModulOmics with subsets of three omics at a time (Figure S9). We found that using all four data sources improves the identification of functionally coherent modules in 92% of the tested cases, with the exception of removing the CoExpression test in GBM. However, in that case, even though the resulting modules were more enriched with known KEGG pathways, they were less enriched with known driver genes (Figure S5).

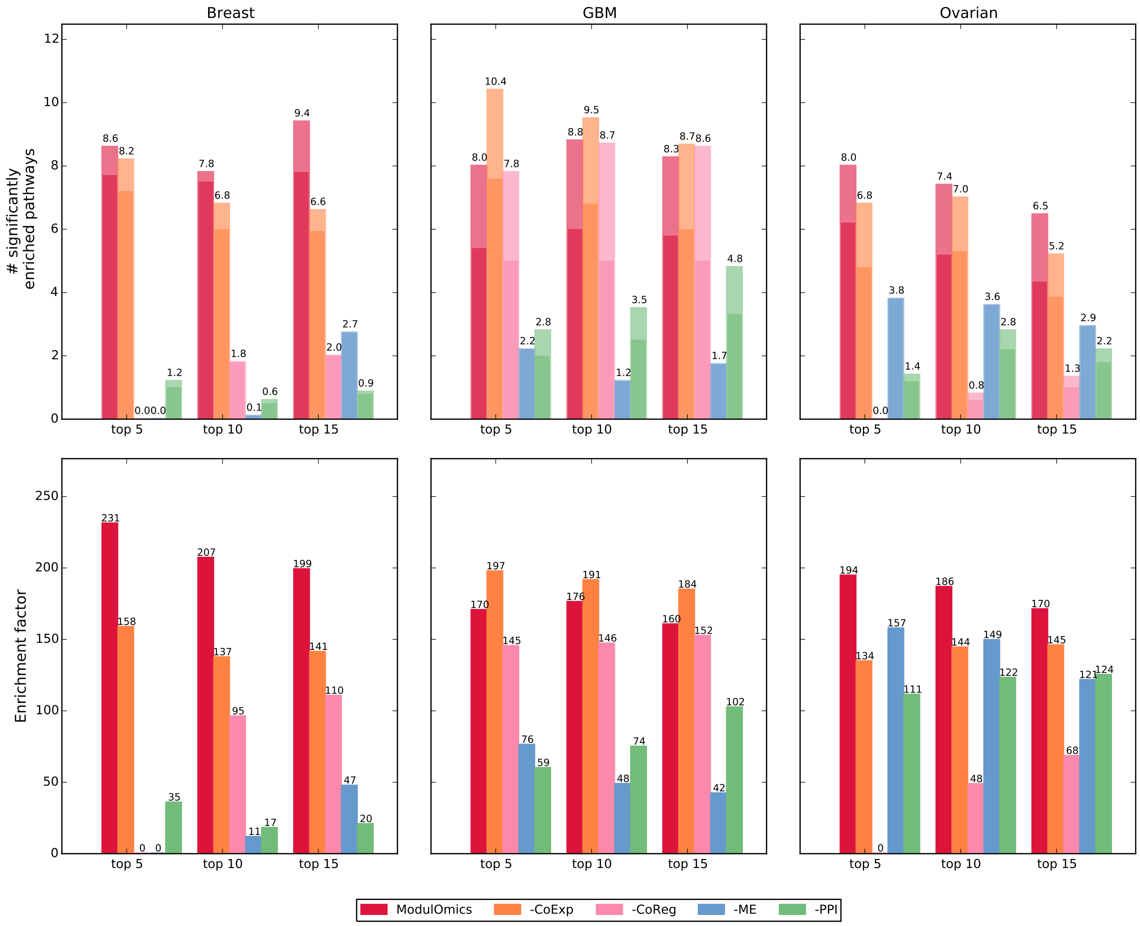

Figure S9: **Pathway enrichment using subsets of three omics.** The − sign next to an omic data source represents the situation in which that omic data source was excluded from the computation of the ModulOmics score.

##### 3.4.3 Module members participating in enriched pathways

Table S7: Genes in top modules participating in enriched pathways (*e.p.*).

| cancer | # top modules | ModulOmics | | | HotNet2 | | | TiMEx | | | MEMCover | | |
|---|---|---|---|---|---|---|---|---|---|---|---|---|---|
| | | # genes in e.p. | # total genes | ratio | # genes in e.p. | # total genes | ratio | # genes in e.p. | # total genes | ratio | # genes in e.p. | # total genes | ratio |
| Breast | 5 | 4 | 5 | 0.8 | 6 | 20 | 0.3 | 2 | 7 | 0.28 | 6 | 12 | 0.5 |
| Breast | 10 | 4 | 7 | 0.57 | 8 | 30 | 0.26 | 2 | 12 | 0.16 | 8 | 23 | 0.34 |
| Breast | 15 | 5 | 10 | 0.5 | 8 | 40 | 0.2 | 3 | 17 | 0.17 | 11 | 35 | 0.31 |
| GBM | 5 | 5 | 6 | 0.83 | 9 | 18 | 0.5 | 0 | 9 | 0.0 | 9 | 11 | 0.81 |
| GBM | 10 | 7 | 9 | 0.77 | 15 | 26 | 0.57 | 0 | 14 | 0.0 | 13 | 21 | 0.61 |
| GBM | 15 | 8 | 11 | 0.72 | 17 | 30 | 0.56 | 0 | 19 | 0.0 | 17 | 31 | 0.54 |
| Ovarian | 5 | 5 | 5 | 1.0 | 3 | 9 | 0.33 | 2 | 7 | 0.28 | 3 | 12 | 0.25 |
| Ovarian | 10 | 11 | 12 | 0.91 | 3 | 9 | 0.33 | 3 | 13 | 0.23 | 6 | 23 | 0.26 |
| Ovarian | 15 | 13 | 15 | 0.86 | 3 | 9 | 0.33 | 3 | 18 | 0.16 | 6 | 33 | 0.18 |

#### 3.5 Extended analysis of breast cancer subtypes

This section extends the Results section in the main text in which we analyzed modules inferred in breast cancer. We first applied ModulOmics on molecularly defined subtypes of breast cancer classified using the mRNA PAM50 classification. The module size distribution (across sizes 2, 3, and 4) in the top 20 modules shows that modules in Luminal B, Her2 and Basal were generally larger (4 genes), while the majority of Luminal A modules consisted of 3 genes (Figure S10A). The scores of the modules ranged from 0.65 to 0.75 (Figure S10B). The sizes of the Luminal A, Luminal B and Her2 modules were correlated to their respective scores, with smaller modules having higher score. The pair modules contained members with very high mutation frequency, such as *TP53* and *GATA3*, and were characterized by high mutual exclusivity scores.

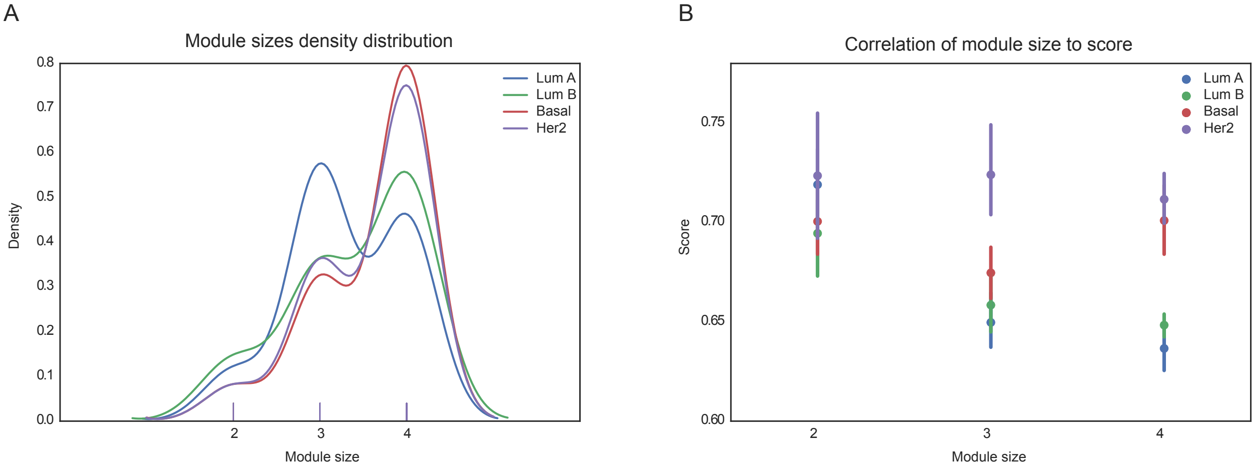

Figure S10: **Size distribution and correlation of size to score** for modules inferred in mRNA-classified breast cancer subtypes, for the top 20 modules of sizes 2, 3, and 4. **A)** Distribution of module sizes (curve smoothed). **B)** Correlation of ModulOmics scores to module size.

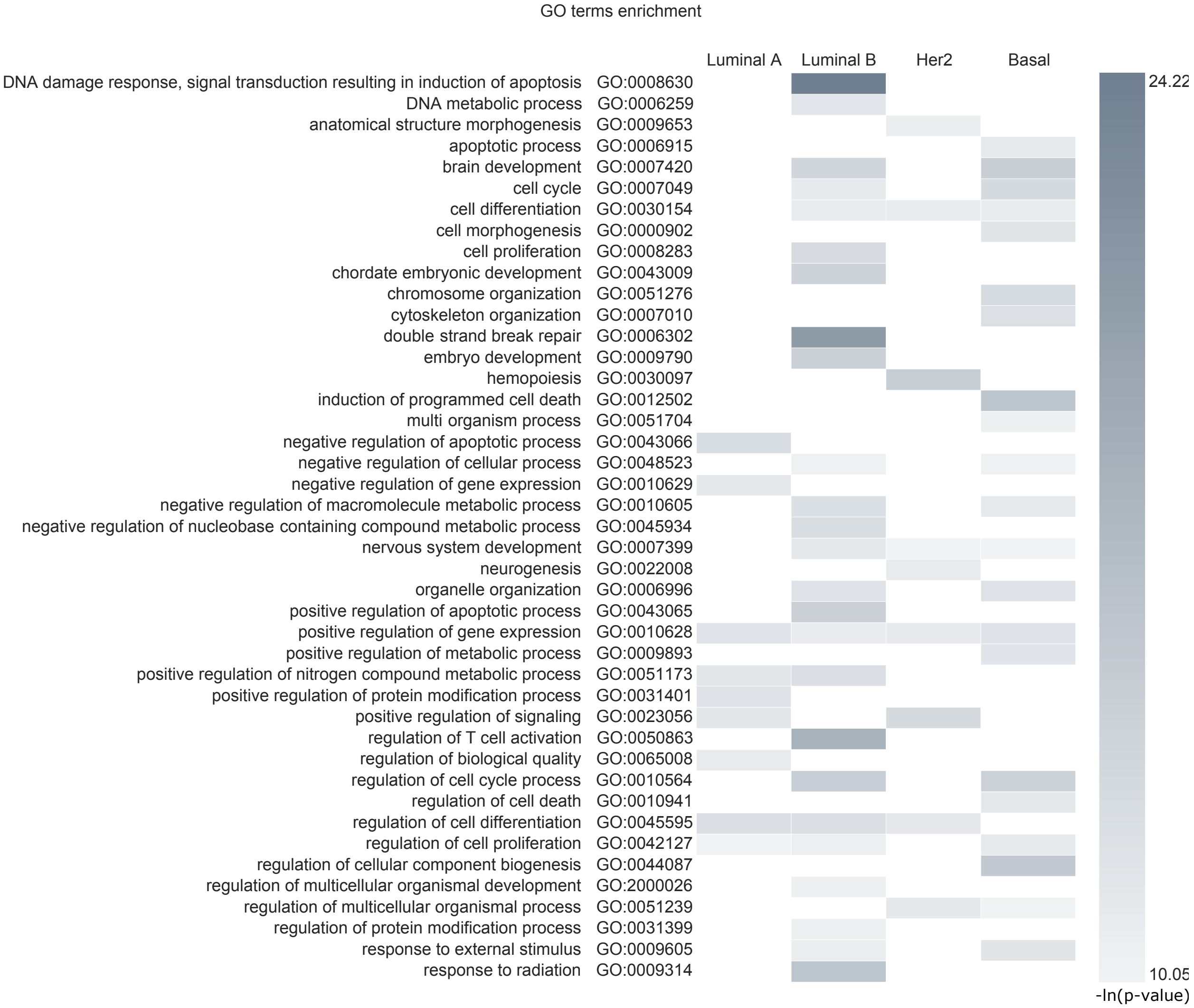

Figure S11: **Significantly enriched *GO* pathways across the top 20 modules (extending Figure 4C in the main text), reflecting the aggressiveness of the Basal and Her2 subtypes, compared to Luminal A and Luminal B**. Enrichment hyper-geometric p-value was computed with Expander [21]. White corresponds to absent pathways.

An alternative way to study breast cancer progression is by stratifying patients according to immunohistochemistry results assessing the HER2, ER and PR receptors. The module size distribution of the top 20 modules was similar to the one obtained by mRNA-based classification (Figure S12A). Modules in Luminal B, Her2-enriched and TN were generally larger (size 4), while the majority of Luminal A modules consisted of 3 genes. We observed a moderate correlation of module size to its score (Figure S12B).

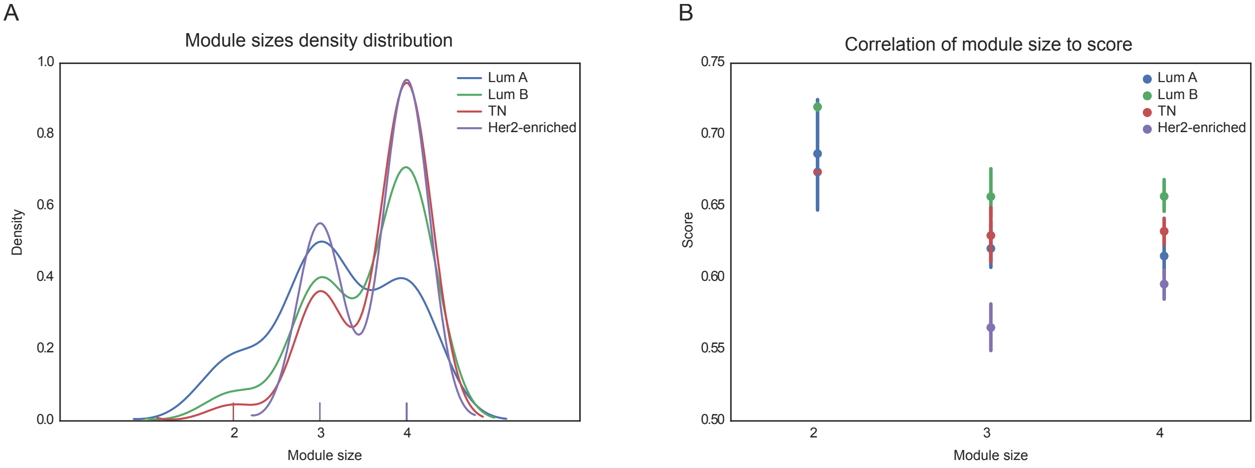

Figure S12: **Size distribution and correlation of size to score** for modules inferred in receptors-classified breast cancer subtypes, for the top 20 modules of sizes 2, 3, and 4. **A)** Distribution of module sizes (curve smoothed). **B** Correlation of ModulOmics scores to module size.

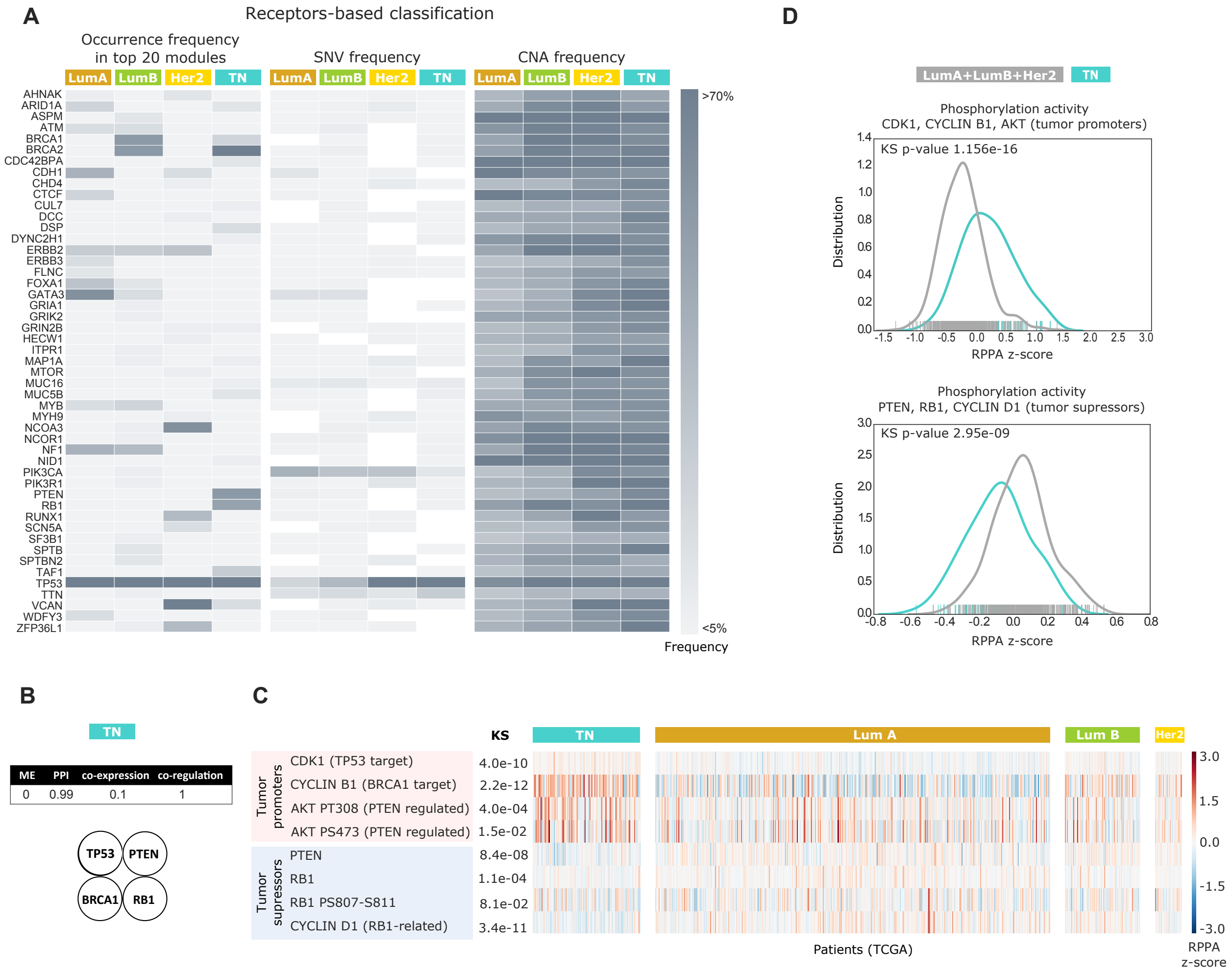

Figure S13: **Modules inferred in receptors-classified breast cancer subtypes highlight differences among subtypes. A)** For each receptors-based subtype and for the pooled set of genes in the top 20 modules, we computed their occurrence frequency in the top 20 modules, as well as their SNV and CNA alteration frequencies across the patient cohort. These genes are enriched with known cancer drivers and pathways, and could not have been identified if relying on SNV and CNA alteration frequencies alone. White corresponds to absent genes (0% frequency). **B)** The TN module *PTEN*, *BRCA1*, *RB1*, and *TP53* (ModulOmics score 0.52) shows both pairwise and group mutual exclusivity scores of 0, unique to this subtype, potentially suggestive that the accumulating activity of multiple tumor suppressors drives TN aggressiveness. **C)** RPPA data supports the observation that TN patients harbor lower activity of the tumor suppressors *PTEN*, *BRCA1*, *RB1*, and *TP53*, compared to other subtypes. PTEN, RB1, phosphorylated RB1 (RB1 PS807-S811) and RB1-related CYCLIN D1 show significantly lower activity, and phosphorylated AKT (AKT PS308 and AKT PS473), which is negatively regulated by *PTEN*, shows higher activity. TP53 and BRCA1 were not captured directly, yet their suppressed targets, CDK1 and CYCLIN B1, show lower activity. **D)** The tumor suppressors *PTEN*, *BRCA1*, *RB1*, and *TP53* can be used to separate TN tissues from the other subtypes in RPPA data. CDK1, CYCLIN B1 and AKT, tumor promoters negatively regulated by *TP53*, *BRCA1* and *PTEN* respectively, show significantly higher activity as a group; PTEN, RB1 and the RB1-related *CYCLIN D1*, tumor suppressors, show significantly lower activity as a group.

Table S8: Top driver modules inferred in mRNA-classified breast cancer subtypes.

| subtype | score | co-expression | co-regulation | ME | PPI | Gene1 | Gene2 | Gene3 | Gene4 |
|---|---|---|---|---|---|---|---|---|---|
| Basal | 0.799 | 0.241 | 1.0 | 1.0 | 0.958 | RB1 | BRCA1 | NF1 | CREBBP |
| Basal | 0.765 | 0.08 | 1.0 | 1.0 | 0.982 | RB1 | BRCA1 | BRCA2 | PTEN |
| Basal | 0.743 | 0.068 | 1.0 | 1.0 | 0.902 | ABCB1 | RB1 | BRCA1 | BRCA2 |
| Basal | 0.741 | 0.263 | 0.75 | 1.0 | 0.95 | RB1 | BRCA1 | BRCA2 | NF1 |
| Basal | 0.741 | 0.08 | 1.0 | 1.0 | 0.884 | ABCB1 | BRCA1 | BRCA2 | PTEN |
| Basal | 0.737 | 0.239 | 0.75 | 1.0 | 0.96 | FMR1 | RB1 | BRCA1 | BRCA2 |
| Basal | 0.731 | 0.267 | 0.75 | 1.0 | 0.906 | RB1 | BRCA1 | BRCA2 | ASPM |
| Basal | 0.726 | 0.017 | 1.0 | 1.0 | 0.887 | ABCB1 | RB1 | BRCA1 | PTEN |
| Basal | 0.718 | 0.175 | 0.75 | 1.0 | 0.948 | APC | BRCA1 | BRCA2 | NF1 |
| Basal | 0.717 | 0.093 | 1.0 | 0.884 | 0.89 | TP53 | BRCA2 | | |
| Basal | 0.712 | 0.211 | 0.75 | 1.0 | 0.887 | BRCA1 | BRCA2 | ASPM | PTEN |
| Basal | 0.703 | 0.139 | 1.0 | 0.703 | 0.97 | TP53 | BRCA1 | BRCA2 | |
| Basal | 0.695 | 0.046 | 1.0 | 0.779 | 0.953 | RB1 | TP53 | BRCA2 | |
| Basal | 0.688 | 0.065 | 1.0 | 0.74 | 0.947 | TP53 | BRCA2 | PTEN | |
| Basal | 0.685 | 0.104 | 1.0 | 0.643 | 0.994 | RB1 | TP53 | BRCA1 | BRCA2 |
| Basal | 0.684 | 0.0 | 1.0 | 0.834 | 0.903 | TP53 | CREBBP | | |
| Basal | 0.679 | 0.0 | 1.0 | 0.772 | 0.945 | TP53 | HSPA4 | CREBBP | |
| Basal | 0.674 | 0.095 | 1.0 | 0.611 | 0.989 | TP53 | BRCA1 | BRCA2 | PTEN |
| Basal | 0.672 | 0.0 | 1.0 | 0.723 | 0.967 | RB1 | TP53 | CREBBP | |
| Basal | 0.67 | 0.0 | 1.0 | 0.694 | 0.988 | RB1 | TP53 | HSPA4 | CREBBP |
| Basal | 0.667 | 0.17 | 0.75 | 1.0 | 0.747 | RB1 | COL12A1 | BRCA1 | NF1 |
| Basal | 0.667 | 0.217 | 0.75 | 0.717 | 0.986 | TP53 | BRCA1 | BRCA2 | HUWE1 |
| Basal | 0.666 | 0.004 | 1.0 | 0.711 | 0.948 | FMR1 | TP53 | HSPA4 | CREBBP |
| Basal | 0.66 | 0.005 | 1.0 | 0.667 | 0.966 | FMR1 | RB1 | TP53 | CREBBP |
| Basal | 0.656 | 0.007 | 1.0 | 0.75 | 0.869 | FMR1 | TP53 | CREBBP | |
| Basal | 0.656 | 0.184 | 0.667 | 0.821 | 0.952 | PRKDC | TP53 | BRCA2 | |
| Basal | 0.653 | 0.108 | 0.75 | 0.778 | 0.975 | WHSC1 | TP53 | BRCA2 | PTEN |
| Basal | 0.649 | 0.123 | 0.75 | 0.747 | 0.978 | TP53 | BRCA2 | HUWE1 | PTEN |
| Basal | 0.648 | 0.106 | 0.75 | 1.0 | 0.737 | DNAH7 | BRCA1 | NF1 | CREBBP |
| Basal | 0.648 | 0.213 | 0.667 | 1.0 | 0.714 | SYNE1 | BRCA1 | NF1 | |
| LumA | 0.737 | 0.1 | 1.0 | 1.0 | 0.847 | TP53 | CDH1 | | |
| LumA | 0.72 | 0.04 | 1.0 | 1.0 | 0.838 | TP53 | FOXA1 | | |
| LumA | 0.703 | 0.259 | 0.667 | 1.0 | 0.888 | TP53 | CDH1 | CTCF | |
| LumA | 0.7 | 0.109 | 1.0 | 1.0 | 0.691 | TP53 | NF1 | | |
| LumA | 0.696 | 0.033 | 1.0 | 0.792 | 0.959 | TP53 | AKT1 | CDH1 | |
| LumA | 0.686 | 0.195 | 0.75 | 0.894 | 0.906 | TP53 | ZFHX3 | CDH1 | CTCF |
| LumA | 0.679 | 0.145 | 1.0 | 0.632 | 0.939 | TP53 | CDH1 | PTEN | |
| LumA | 0.678 | 0.107 | 0.667 | 1.0 | 0.94 | PIK3R1 | AKT1 | PTEN | |
| LumA | 0.665 | 0.192 | 0.75 | 0.752 | 0.964 | TP53 | CDH1 | PTEN | CTCF |
| LumA | 0.659 | 0.202 | 0.667 | 1.0 | 0.768 | TP53 | NF1 | SF3B1 | |
| LumA | 0.651 | 0.13 | 0.75 | 0.75 | 0.974 | TP53 | AKT1 | CDH1 | CTCF |
| LumA | 0.65 | 0.043 | 0.667 | 1.0 | 0.891 | AKT1 | FOXA1 | PTEN | |
| LumA | 0.649 | 0.059 | 0.667 | 1.0 | 0.87 | TP53 | FOXA1 | CTCF | |
| LumA | 0.648 | 0.061 | 0.75 | 0.793 | 0.988 | MAP3K1 | TP53 | AKT1 | CDH1 |
| LumA | 0.642 | 0.243 | 0.667 | 0.809 | 0.849 | TP53 | FOXA1 | GATA3 | |
| LumA | 0.64 | 0.053 | 0.667 | 1.0 | 0.838 | TP53 | FOXA1 | SF3B1 | |
| LumA | 0.639 | 0.149 | 0.75 | 0.7 | 0.958 | TP53 | CDH1 | PTEN | GATA3 |
| LumA | 0.639 | 0.0 | 0.667 | 1.0 | 0.891 | AKT1 | PTEN | MLLT4 | |
| LumA | 0.633 | 0.076 | 0.667 | 1.0 | 0.789 | FOXA1 | PTEN | CTCF | |
| LumA | 0.633 | 0.136 | 0.75 | 0.663 | 0.981 | MAP3K1 | TP53 | CDH1 | PTEN |
| LumA | 0.627 | 0.177 | 0.5 | 0.9 | 0.93 | TP53 | CDH1 | CTCF | SF3B1 |

| | | | | | | | | | |
|---|---|---|---|---|---|---|---|---|---|
| LumA | 0.625 | 0.0 | 0.667 | 1.0 | 0.834 | MED23 | AKT1 | PTEN | |
| LumA | 0.62 | 0.157 | 0.75 | 0.581 | 0.99 | PIK3R1 | TP53 | CDH1 | PTEN |
| LumA | 0.619 | 0.13 | 0.5 | 0.895 | 0.953 | TP53 | CDH1 | MLLT4 | CTCF |
| LumA | 0.618 | 0.166 | 0.667 | 0.807 | 0.832 | MAP3K1 | TP53 | NF1 | |
| LumA | 0.617 | 0.156 | 0.5 | 0.95 | 0.862 | TP53 | CDH1 | CSMD1 | CTCF |
| LumA | 0.617 | 0.316 | 0.667 | 0.71 | 0.775 | FOXA1 | PTEN | GATA3 | |
| LumA | 0.617 | 0.163 | 0.5 | 0.837 | 0.97 | PIK3R1 | TP53 | CDH1 | CTCF |
| LumA | 0.617 | 0.223 | 0.75 | 0.567 | 0.929 | FOXA1 | CDH1 | PTEN | GATA3 |
| LumA | 0.617 | 0.0 | 0.667 | 0.872 | 0.93 | AKT1 | PTEN | PIK3CA | |
| LumB | 0.718 | 0.025 | 1.0 | 1.0 | 0.847 | TP53 | CDH1 | | |
| LumB | 0.718 | 0.008 | 1.0 | 0.904 | 0.959 | TP53 | AKT1 | CDH1 | |
| LumB | 0.702 | 0.066 | 1.0 | 1.0 | 0.743 | CDH1 | GATA3 | | |
| LumB | 0.691 | 0.098 | 1.0 | 1.0 | 0.665 | PTEN | GATA3 | | |
| LumB | 0.687 | 0.037 | 1.0 | 0.771 | 0.939 | TP53 | CDH1 | PTEN | |
| LumB | 0.67 | 0.104 | 0.667 | 1.0 | 0.911 | TP53 | HUWE1 | CDH1 | |
| LumB | 0.668 | 0.179 | 0.75 | 0.812 | 0.931 | AXL | TP53 | CDH1 | PTPRD |
| LumB | 0.667 | 0.0 | 1.0 | 1.0 | 0.67 | ATM | GATA3 | | |
| LumB | 0.664 | 0.087 | 0.75 | 0.873 | 0.945 | AXL | TP53 | HUWE1 | CDH1 |
| LumB | 0.662 | 0.093 | 0.75 | 0.863 | 0.942 | FLNB | AXL | TP53 | CDH1 |
| LumB | 0.661 | 0.047 | 0.75 | 0.876 | 0.973 | TP53 | AKT1 | CDH1 | PTPRD |
| LumB | 0.657 | 0.126 | 0.75 | 0.865 | 0.886 | NF1 | CDH1 | PTEN | GATA3 |
| LumB | 0.654 | 0.146 | 0.5 | 1.0 | 0.971 | TP53 | HUWE1 | CDH1 | ARID1A |
| LumB | 0.653 | 0.067 | 0.75 | 0.828 | 0.968 | FLNB | TP53 | CDH1 | PTEN |
| LumB | 0.652 | 0.115 | 0.667 | 0.904 | 0.924 | TRRAP | TP53 | CDH1 | |
| LumB | 0.649 | 0.111 | 0.75 | 0.782 | 0.953 | AXL | TRRAP | TP53 | CDH1 |
| LumB | 0.649 | 0.021 | 0.75 | 0.852 | 0.975 | FLNB | TP53 | AKT1 | CDH1 |
| LumB | 0.649 | 0.028 | 0.667 | 1.0 | 0.899 | ATM | CDH1 | PTEN | |
| LumB | 0.648 | 0.083 | 0.667 | 1.0 | 0.843 | CDH1 | PTEN | GATA3 | |
| LumB | 0.647 | 0.144 | 0.667 | 1.0 | 0.777 | CDH1 | PTPRD | GATA3 | |
| LumB | 0.646 | 0.127 | 0.667 | 1.0 | 0.79 | HUWE1 | CDH1 | GATA3 | |
| LumB | 0.643 | 0.109 | 0.75 | 0.785 | 0.926 | AXL | TP53 | CDH1 | GATA3 |
| LumB | 0.641 | 0.102 | 0.5 | 1.0 | 0.963 | FLNB | TP53 | CDH1 | ARID1A |
| LumB | 0.64 | 0.135 | 0.75 | 0.715 | 0.96 | TP53 | CDH1 | PTEN | PTPRD |
| LumB | 0.639 | 0.067 | 0.75 | 0.781 | 0.958 | TP53 | CDH1 | PTEN | GATA3 |
| LumB | 0.636 | 0.142 | 0.667 | 1.0 | 0.734 | NF1 | CDH1 | GATA3 | |
| LumB | 0.635 | 0.071 | 0.5 | 1.0 | 0.969 | ERCC6 | AKT1 | CDH1 | ARID1A |
| LumB | 0.632 | 0.022 | 0.75 | 0.803 | 0.952 | MDN1 | TP53 | AKT1 | CDH1 |
| LumB | 0.631 | 0.268 | 0.667 | 0.793 | 0.795 | AXL | TP53 | PTPRD | |
| LumB | 0.626 | 0.091 | 0.75 | 0.696 | 0.966 | AXL | TP53 | CDH1 | PTEN |
| Her2 | 0.81 | 0.387 | 1.0 | 0.925 | 0.928 | XPO1 | TP53 | BRCA2 | |
| Her2 | 0.783 | 0.273 | 1.0 | 0.875 | 0.983 | XPO1 | TP53 | BRCA2 | EGFR |
| Her2 | 0.764 | 0.4 | 0.75 | 0.953 | 0.952 | XPO1 | SMC4 | TP53 | BRCA2 |
| Her2 | 0.757 | 0.141 | 1.0 | 0.925 | 0.96 | TP53 | BRCA2 | EGFR | |
| Her2 | 0.755 | 0.13 | 1.0 | 1.0 | 0.89 | TP53 | BRCA2 | | |
| Her2 | 0.75 | 0.424 | 0.75 | 0.875 | 0.952 | CASC5 | XPO1 | TP53 | BRCA2 |
| Her2 | 0.742 | 0.395 | 0.75 | 0.875 | 0.949 | DSP | XPO1 | TP53 | BRCA2 |
| Her2 | 0.728 | 0.078 | 1.0 | 0.853 | 0.982 | TYK2 | TP53 | EGFR | CDH1 |
| Her2 | 0.723 | 0.18 | 1.0 | 0.769 | 0.942 | JAK2 | TP53 | CDH1 | |
| Her2 | 0.721 | 0.156 | 1.0 | 0.736 | 0.993 | JAK2 | TP53 | EGFR | CDH1 |
| Her2 | 0.72 | 0.377 | 0.75 | 0.781 | 0.971 | DLG1 | XPO1 | TP53 | BRCA2 |
| Her2 | 0.717 | 0.065 | 1.0 | 0.879 | 0.924 | TYK2 | TP53 | EGFR | |
| Her2 | 0.717 | 0.052 | 1.0 | 0.925 | 0.891 | TP53 | BRCA2 | PAX5 | |
| Her2 | 0.707 | 0.212 | 0.75 | 0.892 | 0.976 | XPO1 | TP53 | BRCA2 | CDH1 |
| Her2 | 0.707 | 0.135 | 1.0 | 0.714 | 0.977 | JAK2 | TYK2 | TP53 | CDH1 |

| | | | | | | | | | |
|---|---|---|---|---|---|---|---|---|---|
| Her2 | 0.707 | 0.103 | 1.0 | 0.769 | 0.955 | TP53 | EGFR | CDH1 | |
| Her2 | 0.702 | 0.078 | 1.0 | 0.783 | 0.946 | TP53 | ZFHX3 | EGFR | CDH1 |
| Her2 | 0.702 | 0.001 | 1.0 | 0.908 | 0.898 | TYK2 | TP53 | CDH1 | |
| Her2 | 0.701 | 0.291 | 0.75 | 0.846 | 0.918 | XPO1 | TP53 | BRCA2 | FLG |
| Her2 | 0.701 | 0.129 | 0.75 | 0.946 | 0.978 | TP53 | BRCA2 | MCM6 | EGFR |
| Her2 | 0.696 | 0.302 | 0.667 | 0.925 | 0.89 | CASC5 | TP53 | BRCA2 | |
| Her2 | 0.692 | 0.0 | 1.0 | 1.0 | 0.769 | TYK2 | TP53 | | |
| Her2 | 0.691 | 0.106 | 0.75 | 0.946 | 0.962 | IPO8 | TP53 | BRCA2 | EGFR |
| Her2 | 0.69 | 0.182 | 0.75 | 0.853 | 0.975 | TP53 | EGFR | CDH1 | CUL4B |
| Her2 | 0.689 | 0.224 | 0.75 | 0.819 | 0.963 | XPO1 | TP53 | EGFR | CUL4B |
| Her2 | 0.687 | 0.175 | 0.75 | 0.853 | 0.972 | JAK2 | TP53 | CDH1 | CUL4B |
| Her2 | 0.687 | 0.168 | 0.667 | 1.0 | 0.912 | TP53 | BRCA2 | TBL1XR1 | |
| Her2 | 0.686 | 0.113 | 0.75 | 0.892 | 0.991 | TP53 | BRCA2 | EGFR | CDH1 |
| Her2 | 0.677 | 0.12 | 0.75 | 0.859 | 0.978 | JAK2 | TYK2 | TP53 | BRCA2 |
| Her2 | 0.675 | 0.158 | 0.75 | 0.806 | 0.987 | JAK2 | TP53 | BRCA2 | CDH1 |

Table S9: Top driver modules inferred in receptors-classified breast cancer subtypes. Her2 refers to Her2-enriched.

| subtype | score | co-expression | co-regulation | ME | PPI | Gene1 | Gene2 | Gene3 | Gene4 |
|---|---|---|---|---|---|---|---|---|---|
| TN | 0.708 | 0.118 | 0.75 | 1.0 | 0.964 | MAP1A | RB1 | BRCA1 | BRCA2 |
| TN | 0.706 | 0.127 | 1.0 | 0.807 | 0.89 | TP53 | BRCA2 | | |
| TN | 0.679 | 0.213 | 0.667 | 0.886 | 0.952 | TAF1 | TP53 | BRCA2 | |
| TN | 0.666 | 0.025 | 0.75 | 1.0 | 0.888 | RB1 | BRCA1 | BRCA2 | MUC5B |
| TN | 0.662 | 0.018 | 1.0 | 0.684 | 0.947 | TP53 | BRCA2 | PTEN | |
| TN | 0.659 | 0.0 | 1.0 | 0.684 | 0.953 | RB1 | TP53 | BRCA2 | |
| TN | 0.657 | 0.011 | 0.75 | 1.0 | 0.867 | RB1 | BRCA1 | MUC5B | PTEN |
| TN | 0.653 | 0.104 | 0.75 | 0.772 | 0.986 | TAF1 | TP53 | BRCA2 | PTEN |
| TN | 0.653 | 0.009 | 0.75 | 1.0 | 0.855 | RB1 | DYNC2H1 | BRCA2 | PTEN |
| TN | 0.651 | 0.009 | 1.0 | 0.611 | 0.985 | RB1 | TP53 | BRCA2 | PTEN |
| TN | 0.649 | 0.085 | 0.75 | 0.772 | 0.991 | RB1 | TAF1 | TP53 | BRCA2 |
| TN | 0.632 | 0.297 | 0.5 | 1.0 | 0.732 | NID1 | MUC16 | PTEN | VCAN |
| TN | 0.63 | 0.089 | 0.75 | 0.698 | 0.981 | TP53 | BRCA2 | PTEN | ARID1A |
| TN | 0.629 | 0.009 | 0.75 | 0.772 | 0.984 | TP53 | GRIN2B | BRCA2 | PTEN |
| TN | 0.628 | 0.107 | 0.667 | 0.798 | 0.941 | TP53 | BRCA2 | ARID1A | |
| TN | 0.626 | 0.093 | 0.75 | 0.698 | 0.962 | DSP | TP53 | BRCA2 | PTEN |
| TN | 0.625 | 0.151 | 0.75 | 0.696 | 0.901 | TAF1 | TP53 | BRCA2 | VCAN |
| TN | 0.625 | 0.085 | 0.75 | 0.671 | 0.992 | TAF1 | TP53 | BRCA1 | BRCA2 |
| TN | 0.624 | 0.176 | 0.5 | 0.861 | 0.961 | DSP | TAF1 | TP53 | BRCA2 |
| TN | 0.622 | 0.054 | 0.75 | 0.698 | 0.987 | RB1 | TP53 | BRCA2 | ARID1A |
| TN | 0.621 | 0.051 | 1.0 | 0.53 | 0.902 | RB1 | TP53 | PTEN | VCAN |
| TN | 0.617 | 0.058 | 0.75 | 0.698 | 0.964 | DSP | RB1 | TP53 | BRCA2 |
| TN | 0.615 | 0.074 | 0.75 | 0.653 | 0.984 | TP53 | BRCA2 | PTEN | CUL7 |
| TN | 0.614 | 0.102 | 1.0 | 0.578 | 0.774 | TP53 | PTEN | VCAN | |
| TN | 0.609 | 0.009 | 0.75 | 0.698 | 0.979 | CDC42BPA | TP53 | BRCA2 | PTEN |
| TN | 0.607 | 0.0 | 0.75 | 0.698 | 0.979 | CDC42BPA | RB1 | TP53 | BRCA2 |
| TN | 0.607 | 0.071 | 0.667 | 0.738 | 0.952 | TP53 | BRCA2 | CUL7 | |
| TN | 0.601 | 0.0 | 0.667 | 0.798 | 0.939 | TP53 | GRIN2B | BRCA2 | |
| TN | 0.597 | 0.0 | 0.667 | 0.798 | 0.924 | TP53 | BRCA2 | CDC42BPA | |
| TN | 0.594 | 0.085 | 0.5 | 0.861 | 0.932 | TAF1 | TP53 | SPTB | BRCA2 |
| LumA | 0.726 | 0.211 | 1.0 | 1.0 | 0.692 | ERBB2 | GATA3 | | |
| LumA | 0.725 | 0.0 | 1.0 | 1.0 | 0.9 | TP53 | ATM | | |

| | | | | | | | | |
|---|---|---|---|---|---|---|---|---|
| LumA | 0.722 | 0.0 | 1.0 | 1.0 | 0.89 | TP53 | BRCA2 | |
| LumA | 0.711 | 0.226 | 1.0 | 0.792 | 0.825 | ERBB2 | MYB | GATA3 |
| LumA | 0.674 | 0.206 | 0.75 | 0.847 | 0.894 | FLNC | ERBB2 | MYB | GATA3 |
| LumA | 0.653 | 0.185 | 0.667 | 1.0 | 0.759 | FLNC | ERBB2 | GATA3 | |
| LumA | 0.648 | 0.211 | 0.75 | 0.713 | 0.919 | ERBB2 | MYB | GATA3 | ARID1A |
| LumA | 0.634 | 0.105 | 1.0 | 0.687 | 0.743 | CDH1 | GATA3 | | |
| LumA | 0.633 | 0.14 | 0.667 | 0.888 | 0.839 | ERBB2 | PTEN | GATA3 | |
| LumA | 0.63 | 0.24 | 0.5 | 0.917 | 0.861 | WDFY3 | TP53 | NF1 | ARID1A |
| LumA | 0.627 | 0.095 | 0.5 | 0.947 | 0.965 | TP53 | ATM | CDH1 | CTCF |
| LumA | 0.627 | 0.0 | 1.0 | 0.817 | 0.691 | TP53 | NF1 | | |
| LumA | 0.623 | 0.115 | 0.667 | 0.888 | 0.822 | TP53 | NF1 | ARID1A | |
| LumA | 0.623 | 0.302 | 0.667 | 0.855 | 0.667 | NF1 | FOXA1 | GATA3 | |
| LumA | 0.617 | 0.212 | 0.667 | 0.844 | 0.744 | ERBB2 | NF1 | GATA3 | |
| LumA | 0.616 | 0.212 | 0.667 | 0.881 | 0.706 | WDFY3 | TP53 | NF1 | |
| LumA | 0.612 | 0.07 | 0.667 | 1.0 | 0.713 | ERBB2 | MYO7A | GATA3 | |
| LumA | 0.61 | 0.059 | 0.667 | 0.874 | 0.84 | TP53 | NF1 | ERBB3 | |
| LumA | 0.609 | 0.106 | 0.667 | 0.831 | 0.832 | MAP3K1 | TP53 | NF1 | |
| LumA | 0.606 | 0.146 | 0.5 | 0.867 | 0.912 | TP53 | NF1 | CDH1 | CTCF |
| LumA | 0.604 | 0.27 | 0.5 | 0.809 | 0.837 | WDFY3 | TP53 | NF1 | AHNAK |
| LumA | 0.604 | 0.133 | 0.5 | 0.877 | 0.905 | TP53 | NF1 | CTCF | ARID1A |
| LumA | 0.601 | 0.268 | 0.667 | 0.644 | 0.827 | FOXA1 | CDH1 | GATA3 | |
| LumA | 0.599 | 0.134 | 0.5 | 0.812 | 0.951 | TP53 | FOXA1 | CDH1 | GATA3 |
| LumA | 0.597 | 0.175 | 0.667 | 0.69 | 0.854 | CDH1 | MYB | GATA3 | |
| LumA | 0.594 | 0.116 | 0.5 | 0.823 | 0.937 | TP53 | CDH1 | GATA3 | CTCF |
| LumA | 0.593 | 0.231 | 0.667 | 0.685 | 0.79 | CDH1 | GATA3 | CTCF | |
| LumA | 0.592 | 0.166 | 0.5 | 0.762 | 0.941 | TP53 | ERBB2 | FOXA1 | GATA3 |
| LumA | 0.59 | 0.158 | 0.5 | 0.853 | 0.848 | WDFY3 | TP53 | NF1 | FOXA1 |
| LumA | 0.589 | 0.069 | 0.667 | 0.838 | 0.784 | TP53 | NF1 | CTCF | |
| LumB | 0.725 | 0.14 | 1.0 | 0.791 | 0.97 | TP53 | BRCA1 | BRCA2 | |
| LumB | 0.722 | 0.0 | 1.0 | 1.0 | 0.89 | TP53 | BRCA2 | | |
| LumB | 0.717 | 0.0 | 1.0 | 1.0 | 0.87 | TP53 | PTEN | | |
| LumB | 0.701 | 0.184 | 0.75 | 0.919 | 0.95 | TP53 | ERBB2 | MYB | GATA3 |
| LumB | 0.696 | 0.099 | 0.75 | 1.0 | 0.936 | TP53 | BRCA1 | NF1 | FOXA1 |
| LumB | 0.694 | 0.239 | 0.75 | 0.866 | 0.922 | TP53 | BRCA1 | BRCA2 | ASPM |
| LumB | 0.694 | 0.214 | 0.75 | 0.855 | 0.957 | TP53 | BRCA1 | BRCA2 | NF1 |
| LumB | 0.675 | 0.07 | 0.75 | 0.889 | 0.989 | TP53 | BRCA1 | BRCA2 | NCOR1 |
| LumB | 0.667 | 0.142 | 0.667 | 1.0 | 0.859 | TP53 | BRCA1 | NF1 | |
| LumB | 0.667 | 0.139 | 0.75 | 0.829 | 0.95 | SPTBN2 | TP53 | BRCA1 | BRCA2 |
| LumB | 0.666 | 0.184 | 0.75 | 0.783 | 0.947 | ERBB2 | PIK3CA | MYB | GATA3 |
| LumB | 0.666 | 0.145 | 0.667 | 1.0 | 0.851 | TP53 | BRCA2 | NF1 | |
| LumB | 0.666 | 0.056 | 0.667 | 1.0 | 0.941 | TP53 | BRCA1 | ERBB2 | |
| LumB | 0.659 | 0.032 | 0.667 | 1.0 | 0.938 | TP53 | BRCA1 | FOXA1 | |
| LumB | 0.655 | 0.07 | 0.75 | 0.829 | 0.971 | MAP1A | TP53 | BRCA1 | BRCA2 |
| LumB | 0.655 | 0.184 | 0.75 | 0.792 | 0.896 | ERBB2 | MYB | GATA3 | TTN |
| LumB | 0.649 | 0.07 | 0.75 | 0.783 | 0.994 | TP53 | BRCA1 | BRCA2 | ATM |
| LumB | 0.648 | 0.085 | 0.667 | 1.0 | 0.84 | TP53 | NF1 | ERBB3 | |
| LumB | 0.644 | 0.139 | 0.667 | 0.91 | 0.85 | TP53 | MYB | GATA3 | |
| LumB | 0.641 | 0.108 | 0.5 | 1.0 | 0.957 | PIK3R1 | TP53 | BRCA1 | NF1 |
| LumB | 0.64 | 0.088 | 0.75 | 0.806 | 0.916 | TP53 | BRCA2 | ASPM | ATM |
| LumB | 0.636 | 0.081 | 0.5 | 1.0 | 0.963 | NCOA3 | GRIK2 | BRCA2 | ATM |
| LumB | 0.636 | 0.055 | 0.667 | 1.0 | 0.822 | TP53 | NF1 | ARID1A | |
| LumB | 0.635 | 0.08 | 0.5 | 1.0 | 0.959 | TP53 | ERBB2 | ITPR1 | MYB |
| LumB | 0.633 | 0.123 | 0.5 | 1.0 | 0.909 | TP53 | BRCA2 | GRIA1 | NF1 |
| LumB | 0.632 | 0.091 | 0.5 | 1.0 | 0.937 | TP53 | SPTB | BRCA1 | ERBB2 |

| | | | | | | | | | |
|---|---|---|---|---|---|---|---|---|---|
| LumB | 0.631 | 0.002 | 0.667 | 1.0 | 0.855 | TP53 | ERBB2 | NF1 | |
| LumB | 0.63 | 0.114 | 0.5 | 1.0 | 0.905 | ZFP36L1 | TP53 | BRCA2 | NF1 |
| LumB | 0.626 | 0.0 | 0.667 | 0.879 | 0.959 | TP53 | BRCA2 | ATM | |
| LumB | 0.626 | 0.127 | 0.5 | 1.0 | 0.878 | TP53 | SPTB | BRCA1 | NF1 |
| Her2 | 0.636 | 0.307 | 0.5 | 0.918 | 0.819 | ZFP36L1 | TP53 | RUNX1 | VCAN |
| Her2 | 0.625 | 0.208 | 0.5 | 0.918 | 0.876 | NCOA3 | TP53 | RUNX1 | VCAN |
| Her2 | 0.618 | 0.227 | 0.667 | 0.882 | 0.698 | TP53 | RUNX1 | VCAN | |
| Her2 | 0.614 | 0.165 | 0.5 | 0.918 | 0.873 | TP53 | CDH1 | RUNX1 | VCAN |
| Her2 | 0.613 | 0.18 | 0.5 | 0.918 | 0.853 | NCOA3 | ZFP36L1 | TP53 | VCAN |
| Her2 | 0.611 | 0.097 | 0.5 | 0.878 | 0.969 | NCOA3 | TP53 | SCN5A | ERBB2 |
| Her2 | 0.607 | 0.078 | 0.5 | 0.878 | 0.973 | NCOA3 | TP53 | ERBB2 | RUNX1 |
| Her2 | 0.603 | 0.247 | 0.5 | 0.83 | 0.837 | TP53 | RUNX1 | MYH9 | VCAN |
| Her2 | 0.602 | 0.094 | 0.667 | 0.882 | 0.765 | NCOA3 | TP53 | VCAN | |
| Her2 | 0.602 | 0.129 | 0.5 | 0.918 | 0.861 | NCOA3 | TP53 | SCN5A | VCAN |
| Her2 | 0.6 | 0.058 | 0.5 | 0.878 | 0.963 | NCOA3 | ZFP36L1 | TP53 | ERBB2 |
| Her2 | 0.598 | 0.049 | 0.5 | 0.878 | 0.967 | NCOA3 | MTOR | TP53 | ERBB2 |
| Her2 | 0.597 | 0.3 | 0.5 | 0.759 | 0.829 | TP53 | AHNAK | RUNX1 | VCAN |
| Her2 | 0.593 | 0.007 | 0.5 | 0.878 | 0.987 | NCOA3 | TP53 | GRIN2B | ERBB2 |
| Her2 | 0.593 | 0.165 | 0.667 | 0.882 | 0.659 | ZFP36L1 | TP53 | VCAN | |
| Her2 | 0.592 | 0.101 | 0.5 | 0.918 | 0.849 | ZFP36L1 | TP53 | CDH1 | VCAN |
| Her2 | 0.591 | 0.15 | 0.5 | 0.918 | 0.795 | ZFP36L1 | TP53 | SCN5A | VCAN |
| Her2 | 0.589 | 0.19 | 0.5 | 0.759 | 0.908 | NCOA3 | DCC | TP53 | VCAN |
| Her2 | 0.587 | 0.035 | 0.667 | 0.882 | 0.763 | TP53 | CDH1 | VCAN | |
| Her2 | 0.587 | 0.211 | 0.5 | 0.759 | 0.877 | NCOA3 | TP53 | AHNAK | VCAN |
| Her2 | 0.566 | 0.047 | 0.667 | 0.882 | 0.67 | TP53 | SCN5A | VCAN | |
| Her2 | 0.563 | 0.076 | 0.5 | 0.759 | 0.915 | NCOA3 | TP53 | ERBB2 | VCAN |
| Her2 | 0.557 | 0.143 | 0.5 | 0.759 | 0.824 | HECW1 | TP53 | CDH1 | VCAN |
| Her2 | 0.555 | 0.181 | 0.667 | 0.882 | 0.492 | TP53 | MYH8 | VCAN | |
| Her2 | 0.545 | 0.0 | 0.667 | 0.882 | 0.63 | SYNE1 | TP53 | VCAN | |
| Her2 | 0.543 | 0.0 | 0.667 | 0.882 | 0.624 | SMG1 | TP53 | VCAN | |
| Her2 | 0.54 | 0.062 | 0.667 | 0.882 | 0.552 | TP53 | ASPM | VCAN | |
| Her2 | 0.539 | 0.209 | 0.0 | 1.0 | 0.948 | NCOA3 | ZFP36L1 | RUNX1 | TP53 |
| Her2 | 0.537 | 0.0 | 0.667 | 0.882 | 0.6 | TP53 | WNK1 | VCAN | |
| Her2 | 0.532 | 0.062 | 0.667 | 0.832 | 0.567 | TP53 | HRNR | VCAN | |

##### 3.6 Demonstrating the generality of ModulOmics via complexes detection

We explored a proof-of-concept example for generalizing ModulOmics beyond identifying cancer driver modules. To this end, we collected known biological complexes from CORUM [17], resulting in 432 complexes of size 2, 501 complexes of size 3, and 179 complexes of size 5, and matched these with random groups of genes of the same size. For both the true complexes and the false random groups, we calculated the ModulOmics score using the breast cancer dataset [2], since it represented the largest cohort used in this study. biggest cohort of cancer expression - the breast cancer study, the HIPPIE PPI network and the TRRUST regulatory network, as they were used for detection of cancer modules. Using the *sklearn* Python package, we trained a logistic regression classifier to distinguish between true complexes and random groups, using the sklearn default suggested parameters (L2 regularization with C=1.0 factor to balance overfitting). The resulted classifier distinguished between the instances with an AUPR of 0.84 across all complex sizes S14, and with an AUPR of 0.9 when focusing only on modules of size 4, suggesting that bigger modules are more distinct that randomly generated groups of same size.

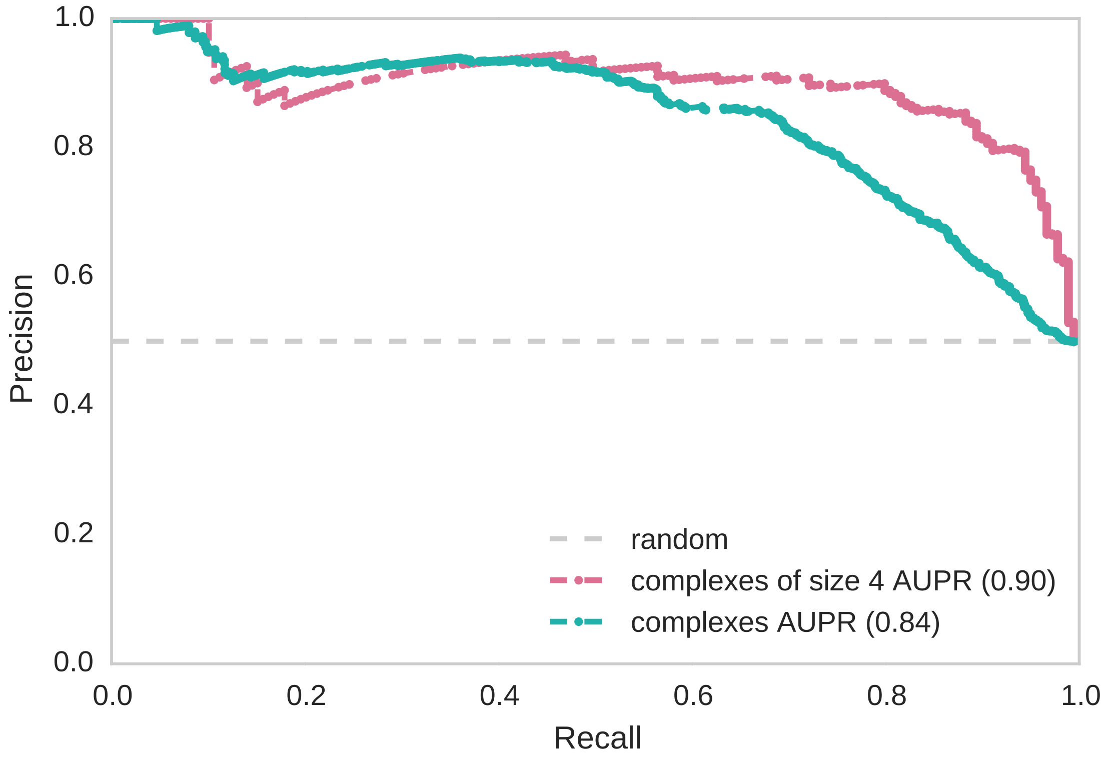

Figure S14: **Precession recall results for classifying known biological complexes and random group of genes**, showing the classification of 1,112 known complexes (blue curve) and a restricted version of 179 known complexes of size 4 (pink curve).

### References

[1] Omer An, Giovanni M. Dall'Olio, Thanos P. Mourikis, and Francesca D. Ciccarelli. Ncg 5.0: updates of a manually curated repository of cancer genes and associated properties from cancer mutational screenings. Nucleic Acids Research, 44(D1):D992–D999, 2016.

[10] Matan Hofree, Hannah Carter, Jason F Kreisberg, Sourav Bandyopadhyay, Paul S Mischel, Stephen Friend, and Trey Ideker. Challenges in identifying cancer genes by analysis of exome sequencing data. Nature Communications, 7, 2016.

[22] UniProt Consortium. UniProt: a hub for protein information. Nucleic acids research, 43(Database issue):D204–D212, January 2015.
